# Multi-environment GWAS analysis for photosynthetic light use efficiency in *Arabidopsis thaliana*

**DOI:** 10.64898/2026.08.03.742429

**Authors:** Thu-Phuong Nguyen, Pádraic J. Flood, Charles Neris Moreira, Nihal Öztolan Erol, Tom P. J. M. Theeuwen, Jeremy Harbinson, Mark G.M. Aarts

## Abstract

Photosynthesis is acknowledged as a potential target to increase crop yield. Improved photosynthesis may be achieved by conventional breeding, exploiting the available natural genetic variation for photosynthesis traits. This approach is challenging for crops due to limitations in high-throughput photosynthesis phenotyping, the highly polygenic nature of photosynthesis, and its strongly dynamic response to environmental changes. Recent advancements in phenomics make accurate and detailed photosynthesis phenotyping more feasible, with the model species *Arabidopsis thaliana* paving the way for applications in crops. In this study, we examined photosynthesis parameters over time in the global Arabidopsis HapMap diversity panel exposed to three conditions: optimal nutrient supply, low phosphorus supply and low nitrogen supply. Combined with two previous studies on photosynthesis in response to low temperature, and to a one-step change in irradiance from low light to high light, five high-quality datasets were systematically analysed using the same approach (with one million-maker set, uni- and multi-variate analyses). Our findings emphasize the genetic complexity of photosynthesis, detecting hundreds of significant quantitative trait loci, only a small number of which are robust, and of which most are condition specific. Robust loci, found in multiple conditions, exemplify those suited for conferring higher all-round photosynthesis, and targets for marker-assisted selection, contributing to environmental resilience, while the multitude of small-effect conditional loci suggest that genomic selection approaches may be more suited to improve crop photosynthesis.

## Introduction

Photosynthesis is a major win-win biological process for humanity, in which CO_2_, the main factor contributing to global warming, is captured from the atmosphere and sequestered in biomass, which for crops provides us with food and feed, and raw materials for industry. Extra photosynthetically captured carbon can also be used to contribute to an improved root system, so improving crop resource use efficiency, or adding to the slowly turning over soil carbon pool. Many efforts have already been made to raise the awareness of the importance of photosynthesis in reducing CO_2_ levels, increasing crop yield to provide food security, and creating an alternative to fossil carbon as a feedstock for the chemical industry and beyond (Long et al., 2006; Zhu et al., 2010; Dodds and Gross, 2007). Photosynthesis can be biotechnologically engineered in various ways to improve its productivity while diminishing its costs in terms of inputs such as fertilizers or water. A major focus in this respect is on the light-use efficiency with which CO_2_ is captured by plants and converted into carbohydrates; this is the aspect of photosynthesis that contributes to crop yield and is a trait that has not so far been substantially increased by selection and breeding (Zhu et al., 2010). Proof-of-principle successes have been demonstrated for tobacco, soybean and wheat, where increases in photosynthetic efficiency were translated into improvements of plant biomass (Kromdijk et al., 2016; Driever et al., 2017, South et al., 2019; de Souza et al., 2022). In tobacco and soybean, photosynthetic efficiency was improved by increasing the speed with which a photoprotective mechanism in photosynthesis (qE) responded to a decrease in light intensity, thus increasing the efficiency of photosynthesis in fluctuating light (Kromdijk et al., 2016; de Souza et al., 2022). In wheat, improving the regeneration rate of Ribulose-1,5-biphosphate (RuBP) in the Calvin cycle by overexpressing sedoheptulose-1,7- bisphosphatase led to an increased rate of CO_2_ fixation (Driever et al., 2017). However improving photosynthesis is not always correlated with increased biomass accumulation. Garcia-Molina et al. (2020) showed that the same approach used to improve photosynthesis in tobacco (Kromdijk et al., 2016) did not result in biomass improvements when used in *Arabidopsis thaliana*. Nevertheless, increasing the conversion efficiency of intercepted irradiance into biomass, a factor still far below its theoretical maximum (Zhu et al., 2010), is one of the most promising traits to develop to further increase crop yield (Theeuwen et al., 2022).

One of the reasons that plant breeding has not substantially improved the conversion efficiency of photosynthesis is that the genetic variation that exists for this trait in crops and their near wild relatives has been poorly explored (Flood et al., 2011; Theeuwen et al., 2022). It is a genetically and physiologically complex quantitative trait that involves multiple genes with multiple alleles, affecting multiple morphological, anatomical, physiological and biochemical and biophysical aspects of plant function, often with a small effect size. This means genetic improvement of the conversion efficiency by breeding is cumbersome, a problem made worse by the challenge of reproducibly phenotyping environmentally sensitive photosynthetic traits over time in large genetic populations in the field (Murchie et al., 2018), but not impossible (McAusland et al., 2020). The conversion efficiency of photosynthesis refers to the fraction of light energy that is converted into the chemical energy of biomass at the point of harvest. In short, photons are absorbed first by photosynthetic pigments, resulting in the formation of excited and thus energy-rich chlorophyll *a* molecules in photosystems I and II. These excited state molecules can either drive photochemistry, and process that takes place in the specialised structures called reaction centres. Alternatively the excited chlorophylls can lose their energy via thermal dissipation processes, or lose their energy via the emission photons in the form of chlorophyll fluorescence (Cf), essentially reversing the excitation process. In the case of photosystem II (PSII) the way the reaction centre works allows these three processes quantified as yields; ΦPSII, ΦNO and ΦNPQ, where ΦPSII is the light-use efficiency for photochemistry; ΦNO is the yield of the constitutive non-photochemical dissipation processes (which includes chlorophyll fluorescence); and ΦNPQ is the yield of the inducible non-photochemical dissipation processes (often dominated by the qE process). The sum of these yields is 1 so they are proportionally distributed: if one yield changes, the rates of other two must also change. As a result, Cf is a sensitive and extremely useful parameter to assess the light-use efficiency of PSII electron transport (ΦPSII) in particular, and the regulation of PSII in general (Baker, 2008; Murchie and Harbinson, 2014; Harbinson, 2018). As ΦPSII is the efficiency of the light-use efficiency for linear electron transport, which drives photosynthetic metabolism, it has become widely used as an index of overall photosynthetic light-use efficient. Cf-based imaging that allows the measurement and imaging of ΦPSII and other chlorophyll fluorescence-derived parameters has been implemented into an automated high-throughput (HTP) phenotyping platform, the Phenovator, that is capable of screening 1440 *A. thaliana* plants multiple times per day for ΦPSII and other parameters that are connected to PSII or can be measured optically (e.g. leaf chlorophyll content) (Flood et al., 2016).

Since the development of HTP phenotyping platforms for photosynthetic and other parameters, quantitative genetic studies exploring natural variation for traits underlying to these parameters have become more and more feasible. At the same time, thanks to the development of whole genome sequencing, high-density genetic maps have been generated based on single nucleotide polymorphisms (SNPs), for large diversity panels consisting of 100s or 1000s accessions of diverse natural genotypes, that are suitable for genome wide association studies (GWAS). GWAS of the genetic variation for ΦPSII under environmentally challenging conditions has already pinpointed allelic variation at specific genes as the causal agents for phenotypic variation in *A. thaliana* (van Rooijen et al., 2017; Prinzenberg et al., 2020). Van Rooijen et al. (2017) showed that allelic variation in the *YELLOW SEEDLING 1* (*YS1*) gene explained part of the natural diversity in photosynthesis acclimation to high irradiance. The allelic variation in a gene encoding for PSII associated protein PSB27 was validated to affect ΦPSII in the cold (Prinzenberg et al., 2020). These GWAS used the *A. thaliana* HapMap population, a diversity panel that consists of 350 natural accessions. After an initial genotype map holding 215,000 SNPs, at an average density of about one SNP per 500 bp (Kim et al., 2007; Li et al., 2010), an imputed SNP marker set has become available that comprises over one million SNPs (Arouisse et al., 2020). This enables the identification of small genomic regions comprising the SNP associated with the observed phenotype, often containing only a small number of candidate genes, for loci identified in GWAS. GWAS performed for the HapMap population using the imputed SNPs data set, with different stringency thresholds (with 1 and 3 million SNPs), revealed a major locus for growth reduction in plants exposed to heat that was not identified when using only the initial 215000 SNP markers (Arouisse et al., 2020). This illustrates the increase in statistical power to detect phenotype-marker associations when using, the imputed SNP marker set.

The power for quantitative trait locus (QTL) detection via GWAS for a single trait (univariate GWAS) can also be increased by the joint analysis of multiple, potentially correlated traits in a multivariate GWAS approach. This may improve the detection of genetic variants whose effects are too small to be detected in univariate analysis (Amos and Laing, 1993; Jiang and Zeng, 1995; Galesloot et al., 2014). For time-series trait data that are generated in a HTP phenotyping platform such as the Phenovator, there is such genetic correlation between traits at different timepoints, and cross-trait covariance (Zhu and Zhang, 2009), which favours a multivariate approach. Moreover, multivariate GWAS can more reliably reveal any common genetic variant that associates with multiple traits, for example in the case of pleiotropy, than can a cross-trait comparison in univariate GWAS (Galesloot et al., 2014). Thoen et al. (2017) successfully identified common underlying genetic factors in a multivariate GWAS of the HapMap population for 11 traits related to plant response to single and combined stresses.

In this study we have investigated the variation in ΦPSII of the HapMap population in the Phenovator HTP phenotyping platform under three conditions: optimal nutrient supply (Optimal), low phosphorus supply (Low-P) and low nitrogen supply (Low-N). An additional population, the Swedish RegMap sub-population is examined under optimal conditions only. Uni- and multivariate GWAS are employed to identify QTL for ΦPSII in each condition using the 1-M imputed genotype data (Arouisse et al., 2020). We have also reanalysed the published ΦPSII data collected for the HapMap in changing temperature (normal to cold temperature, “Cold”, Prinzenberg et al., 2020) and upon a one-step change in irradiance (low light to high light, “LLHL”, van Rooijen et al., 2017) with the same approach. As a result, we have generated an extensive QTL inventory for ΦPSII, which presents the genetic landscape for the photosynthetic light-use efficiency in *A. thaliana*.

## Results

GWAS for the ΦPSII parameter of photosynthesis were carried out based on individual SNP data to obtain a high resolution of identified QTLs. For visualization QTLs were reported based on 25- kb genome windows, which corresponds well with the relatively small regions of LD decay that are common in *A. thaliana* (Kim et al., 2007). A window is considered to be associated with the trait if it contains at least one SNP with a −log_10_(p)≥5.5 (see Material and methods). First, each data set was analysed separately for each of the conditions we used. Thereafter, the results of all five datasets were integrated into one large inventory of QTLs for ΦPSII to identify common QTLs found in more conditions, and specific QTLs found in one condition. We labelled a QTL according to the condition it is found in and the chromosome it is located on, and numbered it according to the order of identification. For each QTL, we calculated a cumulative association score (CAS) by summing the −log_10_(p) values from all GWAS in which the locus exceeded the significance threshold. This score was not intended as a formal statistical measure of significance, but rather as an empirical indicator integrating both the frequency of detection and the strength of association across multiple GWAS (see Material and methods).

### Photosynthetic light-use efficiency under optimal condition

ΦPSII was measured from 11 to 24 days after sowing (DAS) for the HapMap and the Swedish subset of the RegMap collection of accessions grown under optimal conditions with standard nutrition and light intensity. Plants were measured three times a day (morning, noon and afternoon), which generates data for 41 timepoints (Figure 1A). ΦPSII shows a diurnal fluctuation, with the lowest ΦPSII values measured in the afternoon. While ΦPSII initially increases with the growth of the plants, it reaches a maximum as the plants get older. The interquartile range of ΦPSII is from 0.648 to 0.664 at the morning timepoint at 11 DAS to 0.672 to 0.683 at the morning timepoint at 24 DAS (Figure 1A).

**Figure 1.**
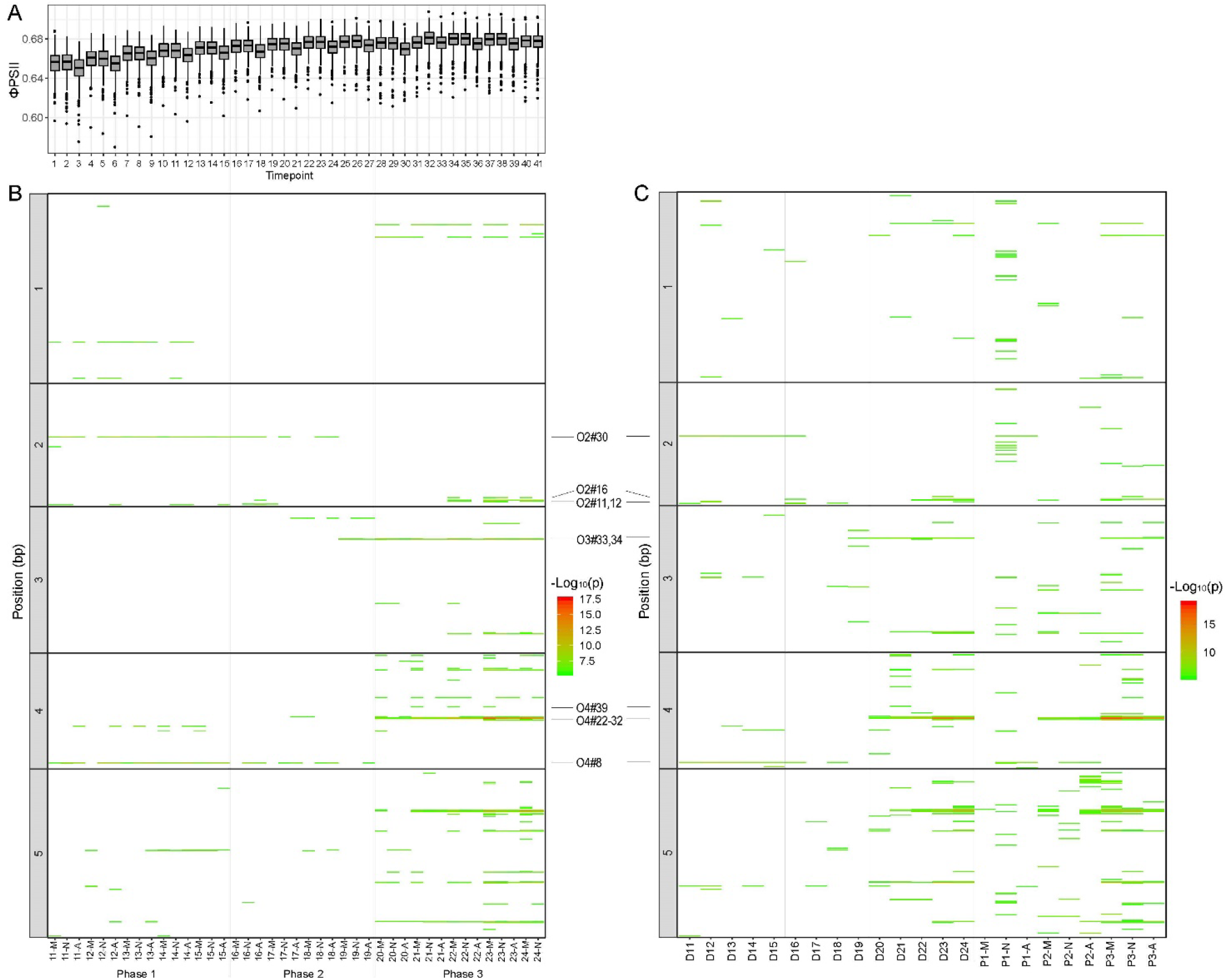
Univariate and multivariate GWAS of photosynthesis (ΦPSII) under optimal conditions. (A) Boxplot of ΦPSII values (y-axis) of the combined set of accessions of the HapMap and Swedish ReqMap panels over time (x-axis): 41 timepoints are shown, based on three measurements per day (morning (M), noon (N), and afternoon (A)) from 11 days after sowing (DAS) till 24 DAS. (B) Univariate GWAS of respective timepoints (x-axis): QTLs are plotted according to their position on the chromosome (y-axis) and significance score (−log_10_(p)) indicated using a colour scale, ranging from green (equal to −log_10_(p)=5.5) to red (equal to a −log_10_(p)=17.5). Three growth phases are distinguished based on their typical QTL profile (phase 1-3). (C) Multivariate GWAS includes Multi-Day GWAS and Multi-Daytime GWAS: in the Multi-Day GWAS model, ΦPSII values of three timepoints of the same day (D) are simultaneously used as input, giving a total of 14 Multi-Day GWAS from 11 to 24 DAS. In the Multi-Daytime GWAS, ΦPSII values of the same daytime-point (M, N or A) in the same growth phase (P) are input traits, giving a total of nine Multi-Daytime GWAS. QTLs are plotted according to their position and significance score on colour scale, ranging from green (−log_10_(p)=5.5) to red (−log_10_(p)=17).

QTLs were identified by univariate GWAS of ΦPSII for each timepoint, and aligned with the phenotypic data of each respective timepoint, which makes up a time-line of QTL profiles (Figure 1B). This reveals three phases during plant growth, each with a typical QTL profile. All QTLs shown in figure 1 have been numbered and listed in the Supplemental Table 1, as there are too many to include in the figure. While many QTLs are only found in one phase, there are also QTLs found in two phases, e.g. QTLs O2#30 and O4#8 in both phase 1 and phase 2 and QTLs O3#33 and O3#34 in both phase 2 and phase 3 (Figure 1B).

Some of the identified QTLs reflect the diurnal response of ΦPSII. An example of this is the QTL region at the top of chromosome 2 (O#11, 12 and 16) which runs from day 22 until the end of the experiment. This QTL is only significant at the morning and noon timepoints but not in the afternoon (Figure 1B). The underlying gene(s) for this QTL are most likely influenced by a diurnal rhythm. QTL O4#8 is constitutively present in the first developmental phase (11 until the end of 15 DAS), except in the morning of 12 DAS, and becomes rhythmic in the second growth phase (significant in the noon and afternoon of 16 to 19 DAS, Figure 1B). These examples suggests that a diurnal regulation of photosynthesis may depend on the plant developmental stage.

Multivariate GWAS was carried out in two ways because of the effect of plant growth and diurnal rhythm on ΦPSII. In the Multi-Day approach, ΦPSII values of three timepoints per day were used as input for the analysis, which results in 14 Multi-Day GWAS outputs for this dataset (Figure 1C). In the Multi-Daytime approach, ΦPSII values measured at the same time each day were used in the analysis. The Multi-Daytime GWAS was performed according to the three growth phases, which leads to nine outputs (three for each timepoint, Figure 1C). The QTL clusters already identified are retained in these multi-GWAS approaches, but further additional QTLs were also found, notably on chromosomes 1, 2 and 3, that were not identified using the univariate GWAS (Supplemental Table 1). Many QTLs are only present in the Multi-Daytime GWAS, further emphasizing the effect of diurnal rhythm on ΦPSII.

In total 64 individual GWAS analyses were performed as a result of combining the uni- and multivariate GWAS for photosynthesis under optimal conditions. The results of this analysis are captured in a QTL inventory shown in Supplemental Figure 2 and Supplemental Table 1. This inventory shows there are in total 331 QTLs were identified, of which 196 are only found in a single GWAS while 135 are found in at least two analyses. QTL O2#30 and O4#8 were the most frequently found significant QTLs were for growth and occurred in phase 1 and 2; they occurred 28 and 32 times with CAS of 189.76 and 232.45, respectively (Supplemental Figure 2, Supplemental Table 1). In growth phase 3, the major QTL O3#33 is significant in 26 times with CAS of 205.59, and the cluster of QTLs on chromosome 4, O4#22-32, is found between 19 to 25 times.

### Photosynthetic light-use efficiency under steady low phosphorus condition

Photosynthetic light-use efficiency has been examined in the HapMap population grown under a constant low P supply. As described above, plant ΦPSII was measured three times a day (morning, noon and afternoon) from 10 to 26 DAS, resulting in 51 timepoints for this dataset. Over the duration of the experiment, ΦPSII shows a consistent diurnal rhythm with a lower ΦPSII being measured in the afternoon timepoints. Similar to what was found for plants grown under optimal conditions, the ΦPSII profile has three phases (Figure 2A). In phase 1, ΦPSII shows a steady increase up to 15 DAS; in phase 2 (16 to 20 DAS) ΦPSII remains stable; and then it decreases slightly during the final six days of phase 3 (21 to 26 DAS). The variation in ΦPSII also changes over the days depending on the phase. In phase 1 the variation is similar between all timepoints of the same day but narrows towards the end of the experiment. For example, the interquartile of ΦPSII among the population is 0.016 (ranging from 0.649 to 0.665) at the morning timepoint 10 DAS, but it reduces to 0.011 at the morning timepoint 15 DAS (ranging from 0.666 to 0.676). Thereafter, the variation varies from day to day, with largest variation at the afternoon timepoint. In general, variation steadily increases during phase 2, with a ΦPSII interquartile at the afternoon timepoint 20 DAS of 0.015. During phase 3, the largest variation is again observed at the afternoon timepoint (interquartile of around 0.02), with little variation between days.

**Figure 2.**
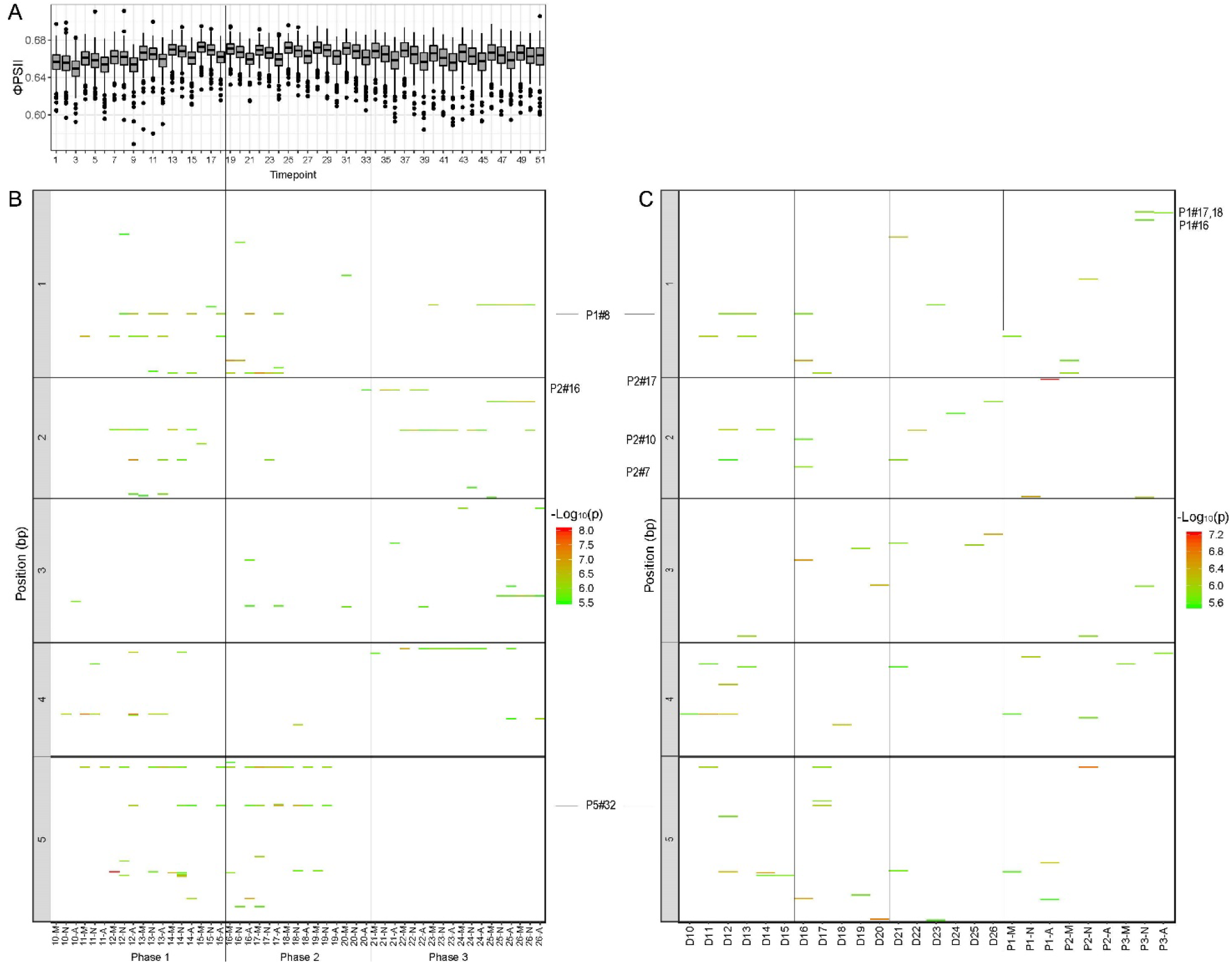
Univariate and multivariate GWAS of photosynthesis (ΦPSII) under low phosphorus supply condition. (A) Boxplot of ΦPSII values (y-axis) of accessions of the HapMap panels over time (x-axis): 51 timepoints are shown, based on three measurements per day (morning (M), noon (N), and afternoon (A)) from 10 days after sowing (DAS) till 26 DAS. (B) Univariate GWAS of respective timepoints (x-axis): QTLs are plotted according to their position on the chromosome (y-axis) and significance score (−log_10_(p)) indicated using a colour scale, ranging from green (equal to −log_10_(p)=5.5) to red (equal to −log_10_(p)=8). (C) Multivariate GWAS includes Multi-Day GWAS and Multi-Daytime GWAS: in Multi-Day GWAS model, ΦPSII values of three timepoints of the same day (D) are simultaneously used as input, giving a total of 17 Multi- Day GWAS from 10 to 26 DAS. In Multi-Daytime GWAS, ΦPSII values of the same daytime-point (M, N or A) and in the same developmental phase (P) are input traits, giving a total of nine Multi-Daytime GWAS. QTLs are plotted according to their position and significance score using a colour scale, ranging from green (−log_10_(p)=5.5) to red (−log_10_(p)=7.2).

Univariate GWAS was performed for ΦPSII measured at the 51 individual timepoints, which results in QTL profiles reflecting the three above mentioned photosynthesis phases (Figure 2B). Twelve QTLs, spread over chromosomes 1, 2, 4 and 5, are repeatedly identified in phase 1, of which five are also identified in phase 2. In total, seven QTLs, locating on chromosomes 1, 3 and 5, are significant in more than one timepoint in phase 2. During phase 3, 13 QTLs are detected, of which eight were also identified in phase 1 and 2, and five are specific for this phase. All the QTLs are numbered (Supplement Table 2), with only those explicitly referred to in the text are indicated in Figure 2B-C.

Only two ΦPSII QTLs under low P supply are affected by the diurnal rhythm. QTL P1#8 first appears in the noon and afternoon measurements of 12 and 13 DAS, but is thereafter only detected in the afternoon from 14 to 17 DAS (Figure 2B). The first significant appearance of QTL P2#16 is in the afternoon at 20 DAS, and in the two days thereafter it is also significant for both noon and afternoon timepoints (Figure 2B).

Similar to the experiment under optimal conditions, multivariate GWAS were performed, of which 17 were Multi-Day GWAS from 10 DAS onwards and 9 were Multi-Daytime GWAS for the morning, noon and afternoon timepoints of the three growth phases (Figure 2C). Only a small number of additional QTLs were detected via the multivariate GWAS, for example, two QTLs on chromosome 2, P2#7 and P2#10, were found in the Multi-Day GWAS for day 16. Multi-Daytime GWAS also resulted in a few additional QTLs: a cluster of QTLs, P1#16, #17 and #18, for phase 3 in the noon and afternoon, and QTL P2#17 for phase 1 in the afternoon.

From a total of 77 individual analyses, 99 QTLs were identified and integrated in an QTL inventory for ΦPSII under low P supply (Supplemental Figure 3, Supplemental Table 2). QTL P5#32 is the most frequently detected QTL (19 times) with CAS of 116.66, and was most often found to be significant (19 times). Chromosomes 1, 2 and 4 also contain several QTLs that were identified at least seven times, with CAS of above 40.

### Photosynthetic light-use efficiency under low nitrogen supply

The photosynthetic light-use efficiency (ΦPSII), of the HapMap set of accessions in response to low N supply, was examined in a similar way for the response to low P supply. ΦPSII was measured three times a day (morning, noon and afternoon), from 12 until 22 DAS, generating 32 data-timepoints (Figure 3A). There was a steady increase of ΦPSII over time, with the median around 0.69, at the first timepoint, to 0.715 at the last timepoint. The phenotypic variation decreases slightly as the plants grew older, with the interquartile of phenotypic variation being around 0.0125 at the first timepoint and 0.010 at last. Diurnal variation in ΦPSII was also observed, with the afternoon timepoint giving the lowest ΦPSII value of the day (Figure 3A).

**Figure 3.**
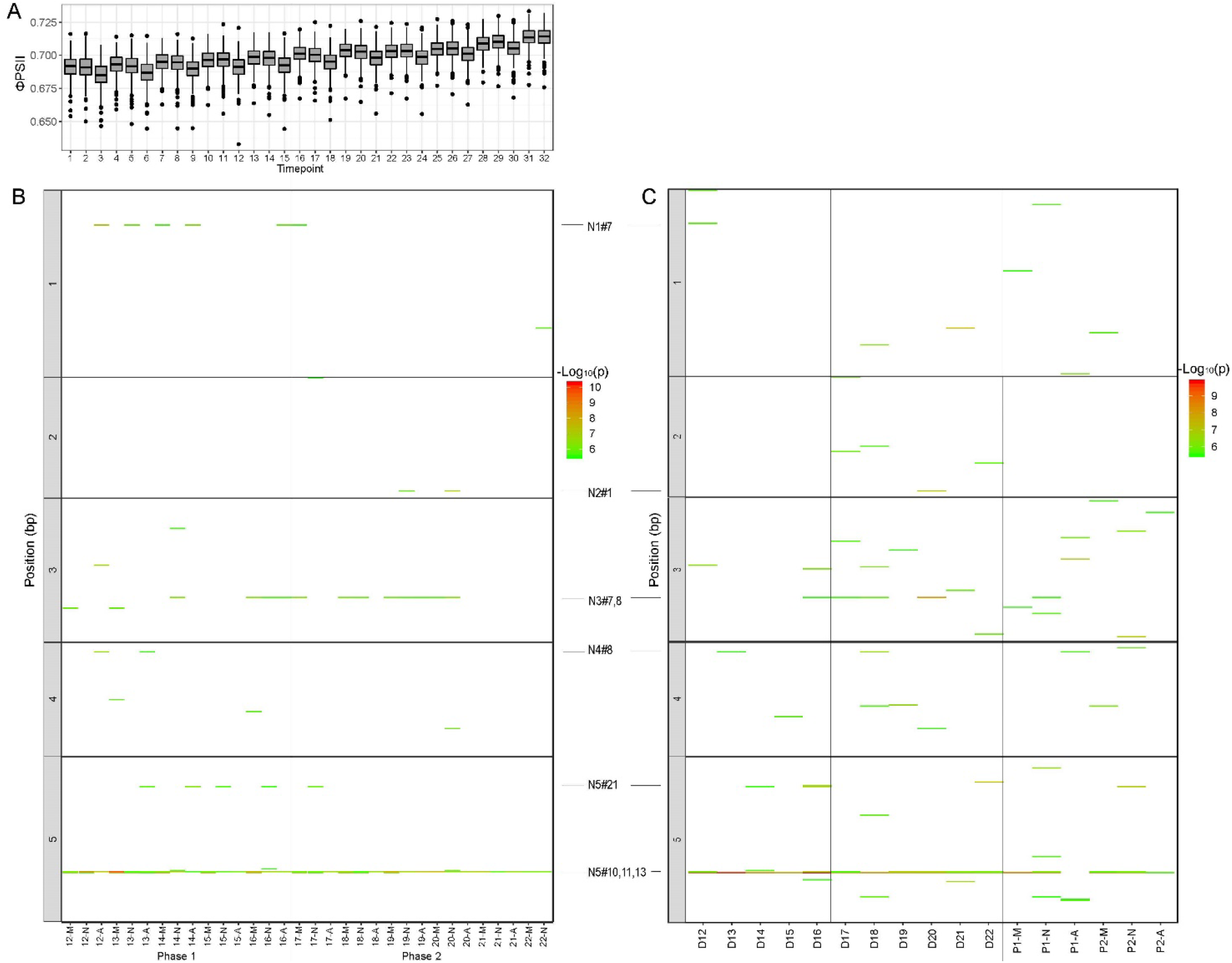
Univariate and multivariate GWAS of photosynthesis (ΦPSII) under low nitrogen supply condition. (A) Boxplot of ΦPSII values (y-axis) of accessions of the HapMap panels over time (x-axis): 32 timepoints are shown, based on three measurements per day (morning (M), noon (N), and afternoon (A)) from 12 days after sowing (DAS) till 22 DAS. (B) Univariate GWAS of respective timepoints (x-axis): QTLs are plotted according to their position on the chromosome (y-axis) and significance score (−log_10_(p)) indicated using a colour scale, ranging from green (equal to −log_10_(p)=5.5) to red (equal to −log_10_(p)=10.2). (C) Multivariate GWAS includes Multi-Day GWAS and Multi-Daytime GWAS: in Multi-Day GWAS model, ΦPSII values of three timepoints of the same day (D) are simultaneously used as input, giving a total of 11 Multi- Day GWAS from 12 to 22 DAS. In Multi-Daytime GWAS, ΦPSII values of the same daytime-point (M, N or A) in the same developmental phase (P) are input traits, giving a total of six Multi-Daytime GWAS. QTLs are plotted according to their position and significance score using a colour scale, ranging from green (−log_10_(p)=5.5) to red (−log_10_(p)=9.8).

The univariate GWAS analyses for the respective timepoints result in 16 QTLs, distributed over all chromosomes. A prominent cluster of three QTLs on chromosome 5, N5#10, #11 and #13, affects photosynthesis under low N supply throughout the whole experiment. Besides this cluster, there are four other major QTLs that are detected multiple times, N1#7, N3#7, N3#8 and N5#21 (Figure 3B). Although ΦPSII shows diurnal variation, only some QTLs reflect a diurnal rhythm, and then only for two days. For example, QTL N2#1 was present at noon at 19 and 20 DAS, and QTL N4#8 was found in the afternoon at 12 and 13 DAS (Figure 3B).

To allow Multi-Daytime GWAS the time-series was divided into two phases with the first phase lasting from 12 to 16 DAS, and second phase lasting from 17 to 22 DAS (Figure 3B). This allows six Multi-Daytime GWAS, one for each timepoint per day, in each phase (Figure 3C). Multi-Day GWAS was also been performed by combining the ΦPSII measurements at all three timepoints of a day, which gives 11 analyses (Figure 3C). In this multivariate approach, we identify many additional QTLs compared to the univariate analysis (Figure 3C), which suggests that there are many common genetic factors affecting ΦPSII at many timepoints though each is likely to have only a small effect size that would not yield a significant association in the univariate analyses. All QTLs are numbered and listed in Supplemental Table 3, and only the ones referred to in the text are indicated in figure 3.

For this dataset, a total of 74 QTLs is found in 49 individual GWAS, of which five QTLs are present in at least 10 analyses (Supplemental Figure 4, Supplemental Table 3): three on chromosome 5, N5#11, #13 and #10, and two on chromosome 3, N3#7 and N3#8. The QTL N5#11, corresponding to the window at 8175000 bp, is found in 48 of the 49 analyses, with CAS of 365.6.

### Photosynthetic light-use efficiency in the cold and upon prolonged darkness

The effect of low temperature (5°C) on ΦPSII in the HapMap population has previously been studied by Prinzenberg et al. (2020). Plants were first grown for 13 days under optimal conditions with a day time temperature of 21°C, then from 14 DAS subjected to cold (5°C, day/night) for one week. On 21 DAS, plants were allowed to recover from the cold for two days at optimal conditions. In this first part of the experiment (part 1) where the effect of changing temperature was examined (Prinzenberg et al., 2020), ΦPSII was measured four times a day, at morning, noon and two afternoon timepoints (Figure 4A). In part 2 of this experiment, plants were exposed to a period of darkness (from 23 to 26 DAS), during which ΦPSII could not be measured. Darkness is known to induce leaf senescence and thus affect photosynthesis (Keech et al., 2007). Plants were allowed to recover from continuous darkness, under optimal temperatures, until the end of the experiment (from 27 to 33 DAS). Variation for ΦPSII at growth irradiance was measured during recovery at four timepoints per day (Figure 4A).

**Figure 4.**
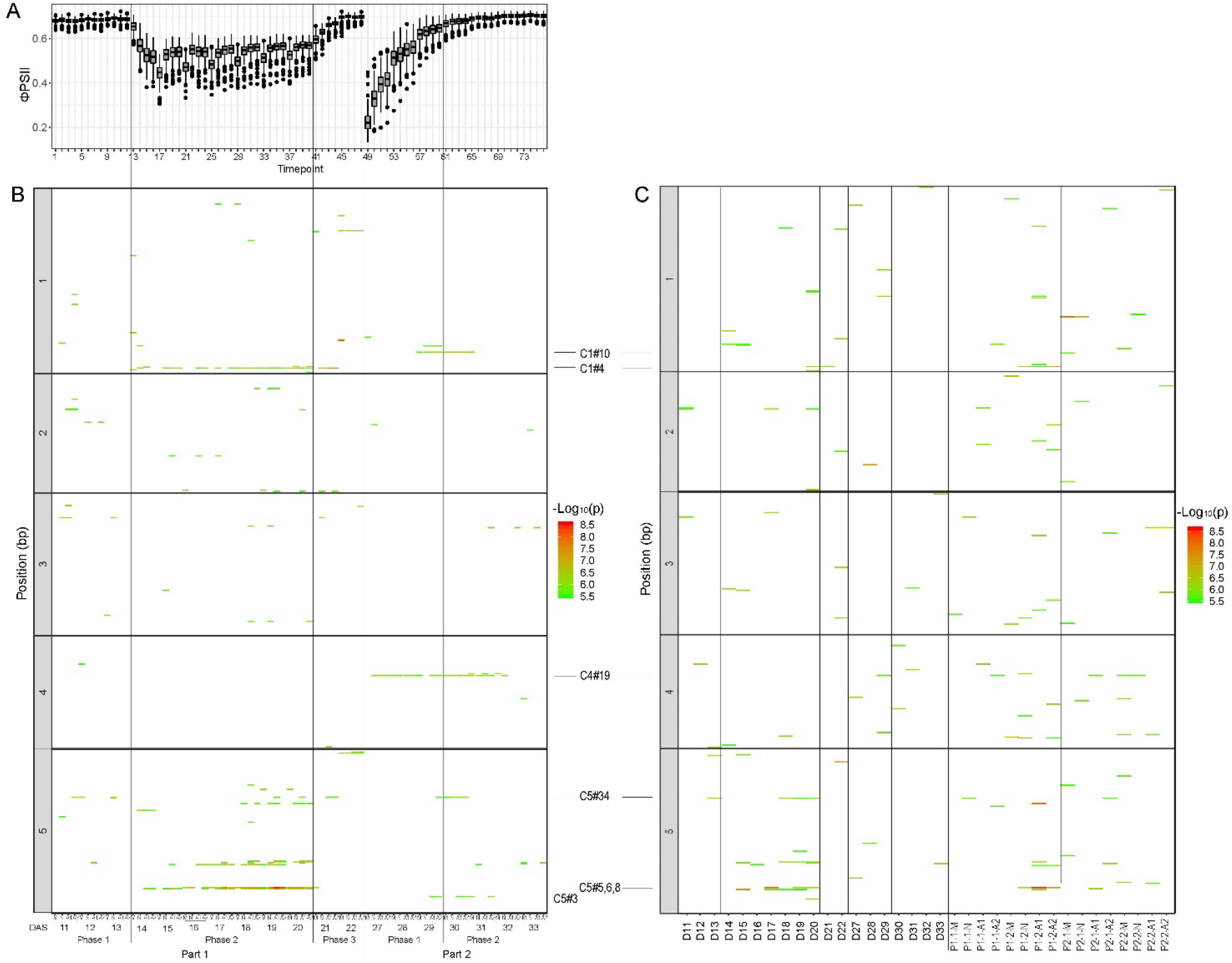
Univariate and multivariate GWAS of photosynthesis (ΦPSII) under changing temperature (from normal to cold, Part 1) and after darkness treatment (Part 2). (A) Boxplot of ΦPSII values (y-axis) of accessions of the HapMap panels over time (x-axis): 76 timepoints are shown, based on four measurements per day (morning (M), noon (N) and 2 times afternoon (A1 and A2)) from 11 days after sowing (DAS) till 22 DAS in part 1, and from 27 to 33 DAS in part 2 of the experiment. (B) Univariate GWAS of respective timepoints (x-axis): QTLs are plotted according to their position on the chromosome (y-axis) and significance score (−log_10_(p)) using a colour scale, ranging from green (equal to −log_10_(p)=5.5) to red (equal to −log_10_(p)=8.6). (C) Multivariate GWAS includes Multi-Day GWAS and Multi-Daytime GWAS: In Multi-Day GWAS model, ΦPSII values of four timepoints of the same day are simultaneously used as input for multivariate model. As a result there are 19 Multi-Day GWAS, from Day 11 (D11) to D22 and from D27 to D33. In Multi-Daytime GWAS, ΦPSII values of the same daytime-point (morning, M, noon, N, and afternoon, A1 or A2) in the same phase (P) according to treatment and response are input traits for multivariate model. There are in total 16 Multi-Daytime GWAS. QTLs are plotted according to their position on the chromosome (y-axis) and significance score using a colour scale, ranging from green (−log_10_(p)=5.5) to red (−log_10_(p)=8.6).

Prinzenberg et al. (2020) reported that upon the exposure of plants to cold, ΦPSII drops substantially in the first day of cold treatment (14 DAS), reaching its the lowest value on the second day of the cold treatment (15 DAS). It then remains low throughout the cold treatment, with a slight increase towards the last day of cold treatment (20 DAS). The diurnal rhythm of the ΦPSII in the population is maintained during the cold treatment with the lowest ΦPSII values in the morning (Figure 4A). The variation of ΦPSII increases strongly upon the start of the cold treatment, but then decreases gradually throughout the treatment. For example, the ΦPSII interquartile ranges from 0.057 at the end of 14 DAS to 0.030 at the end of 20 DAS, while at the regular temperatures, the variation was only around 0.010 (Figure 4A). After four days of darkness, it takes four days for ΦPSII to return to regular, pre-stress levels (Figure 4A). As with the cold treatment, darkness induces additional ΦPSII variation, which decreases gradually to regular levels when plants resume their photosynthetic capacity (Figure 4A). The diurnal variation in ΦPSII is minimal or absent after the exposure to continuous darkness.

Univariate GWAS was performed for each individual timepoint of the experiment, resulting in a total of 76 analyses (Figure 4B). Compared to the previous analysis by Prinzenberg et al. (2020), we used a much larger SNP data set with 1 M SNPs, and set a higher −log_10_(p) threshold of 5.5 for a significant association (compared to previously reported threshold of 3.0). This resulted in us identifying fewer QTLs than Prinzenberg et al (2020), though some of the major QTLs identified previously were found again in this study, such as QTLs #1 (C1#4), #70 (cluster of C5#5-8) and #87 (C5#13). The response of photosynthesis to prolonged darkness (part 2, phase 1) is affected by four major QTLs (C1#10, C4#19, C5#3 and C5#34) (Figure 4B). Only one of those, QTL C5#34, was also occasionally identified during part 1. The little overlap in QTLs found in both treatments means that photosynthesis acclimation after darkness is genetically distinct from response and recovery from cold, and thus likely involves different physiological processes.

Multivariate GWAS was also performed for this dataset including Multi-Day (19 times) and Multi- Daytime (16 times) analyses. Multi-Day analyses identified a number of additional QTLs, especially for part 2 (Figure 4C). Also the Multi-Daytime analyses resulted in several additional QTLs for all treatments and phases (Figure 4C, Supplemental Table 4). Notably, the diurnal response of ΦPSII during the cold period leads to six additional QTLs mapping to chromosome 4, where there was no QTLs found using the univariate GWAS.

A total of 111 GWAS analyses were performed for this experiment, resulting in 170 QTLs (Supplemental Figure 12). The QTL C1#4 (equivalent to QTL#1, and co-locating with the *PSB27* gene identified by Prinzenberg et al. (2020)) is most frequently identified (28 times) with CAS of 181.31. Six other QTLs are found in at least 10 analyses, five on chromosome 5 and one on chromosome 4 (Supplemental Table 4 and Supplemental Figure 5). The QTL on chromosome 4 (C4#19) was specific for photosynthesis efficiency (ΦPSII) during the recovery after darkness, in part 2 of the experiment.

### Photosynthetic light-use efficiency under fluctuating irradiance

Natural variation for ΦPSII in the HapMap population in response to a single step increase in irradiance, from low light (100 μmol m^-2^ s^-1^) to high light (550 μmol m^-2^ s^-1^) (LLHL treatment) was previously studied by van Rooijen et al. (2017). They measured ΦPSII three times a day (morning, noon and afternoon) for two days at low light (23 and 24 DAS, phase 1) and for four days after the switch to high light (25 to 28 DAS, phase 2). For this dataset, we revisited the GWAS analysis, employing the imputed 1M SNP dataset (Arouisse et al., 2020) and applying a higher −log_10_(p)=5.5 threshold for significance (compared to previously reported threshold of 4.0). Furthermore we performed additional multivariate GWAS analyses. The response to the increased irradiance results in a large drop in ΦPSII from around ΦPSII=0.65 at 100 μmol m^-2^ s^-1^ to ΦPSII=0.35 at the first measurement made at 550 μmol m^-2^ s^-1^, after which ΦPSII gradually increases as plants acclimate to the higher irradiance (Figure 5A). Throughout the experiment, ΦPSII displays no diurnal rhythm.

**Figure 5.**
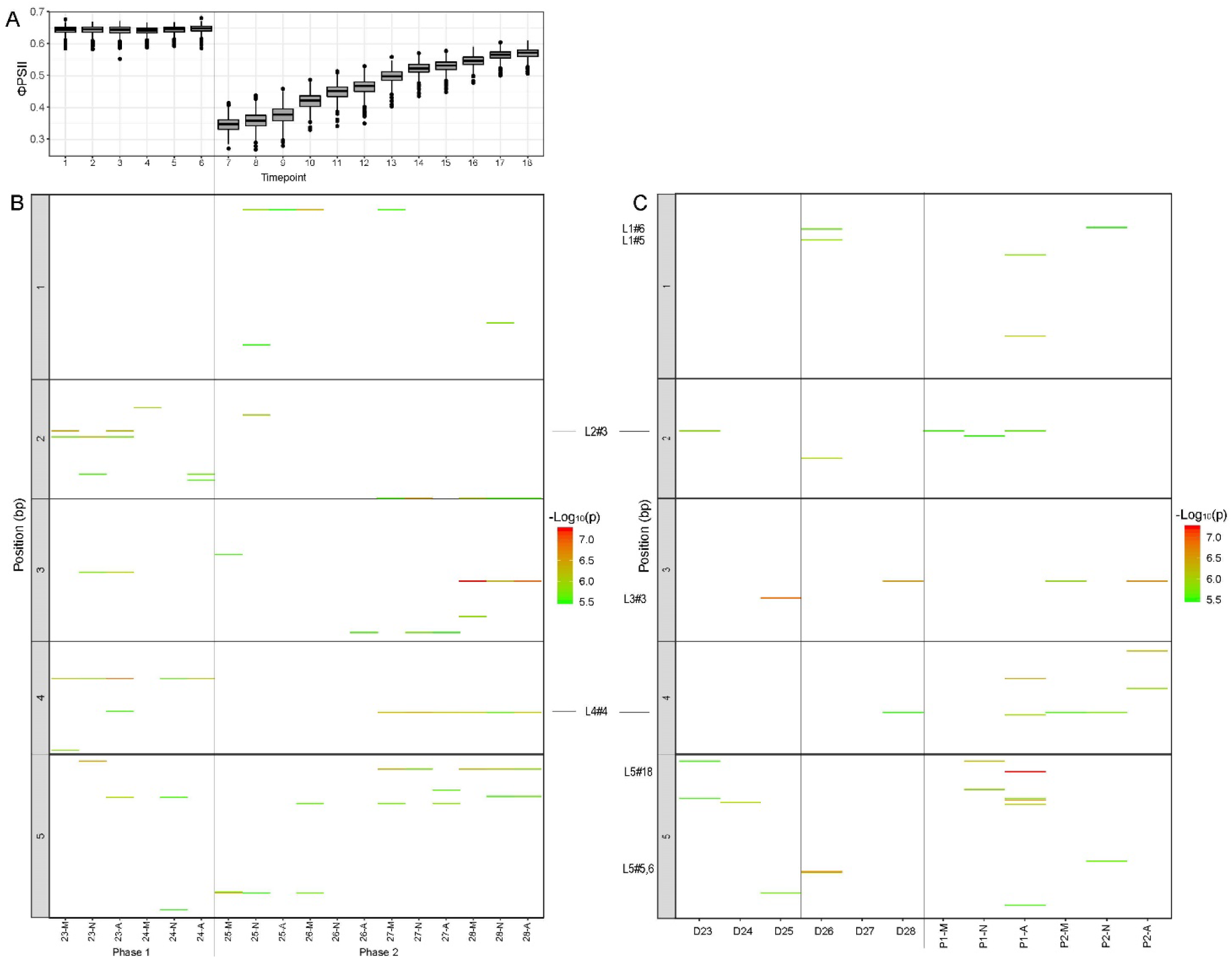
Univariate and multivariate GWAS of photosynthesis (ΦPSII) under changing irradiance (low light to high light). (A) Boxplot of ΦPSII values (y-axis) of accessions of the HapMap panels over time (x-axis): 18 timepoints are shown, based on three timepoints per day (morning (M), noon (N), and afternoon (A)) from 23 days after sowing (DAS) till 28 DAS. (B) Univariate GWAS of respected timepoints (x-axis): QTLs are plotted according to their position on the chromosome (y-axis) and significance score (−log_10_(p)) using a colour scale, ranging from green (equal to −log_10_(p)=5.5) to red (equal to −log_10_(p)=7.5). (C) Multivariate GWAS includes Multi-Day GWAS and Multi-Daytime GWAS: in Multi-Day GWAS model, ΦPSII values of three timepoints of the same day (D) are simultaneously used as input, giving a of total 6 Multi-Day GWAS from 23 to 28 DAS. In Multi-Daytime GWAS, ΦPSII values of the same daytime-point (M, N or A) in the same treatment phase (P) are input traits, giving a total of six Multi-Daytime GWAS. QTLs are plotted according to their position and significance score using a colour scale, ranging from green (−log_10_(p)=5.5) to red (− log_10_(p)=7.5).

Because of the higher significance threshold we now applied, substantially fewer QTLs were identified in the univariate GWAS compared to the original study of van Rooijen et al. (2017). There were no QTLs detected for variation in ΦPSII in both low light and high light (Figure 5B and Supplemental Table 5) suggesting that ΦPSII in low light and in high light are controlled by different physiological mechanisms, involving different genetic factors for which there is variation. Only a few more QTLs with a strong association are identified by the multivariate approach (Figure 5C). For example, Multi-Day GWAS identifies QTL L3#3 (for 25 DAS) or QTL L5#5 and 6 (for 26 DAS), that is not significant in any univariate GWAS for that day (Figure 4C). Although a diurnal rhythm of ΦPSII is not noticed, Multi-Daytime GWAS identifies a new, strong QTL L5#18 for ΦPSII in the afternoon of phase 1 (P1-A, Figure 4C).

The integration of 18 univariate and 12 multivariate analyses results in a total of 54 QTLs (Supplemental Figure 6 and Supplemental Table 5). The most prevalent QTL, L4#4, is found nine times, and has a CSA of 53.81. Ten other QTLs occur at least three times and eight QTLs occur twice, while the others only occur once (Supplemental Figure 6 and Supplemental Table 5).

### Comparison of all QTL inventories for photosynthetic light-use efficiency in multiple environmental conditions

We employed 332 GWAS analyses to identify QTLs for ΦPSII in five datasets obtained from as many independent experiments (Figures 1-5). This resulted in a total of 665 QTL windows (Figure 6A and Supplemental Table 6). We then compared these QTL inventories to identify genetic loci that are common or unique among experiments. Of the 665 QTLs, there are 610 loci that are only found in a single experiment: 294 in Optimal, 71 in the Low P treatment, 62 in the Low N treatment, 140 in the Cold treatment, and 43 in LLHL experiment (Supplemental Table 6). 48 QTLs are found in two experiments and six QTLS are found in three experiments (Figure 6B, Table 1). There is only one QTL found in four out of five experiments (not in the LLHL treatment), mapping to chromosome 5 (window 8275000 bp), equivalent to O5#35, P5#17, N5#13 and C5#18 (Figure 6, Table 1).

**Figure 6.**
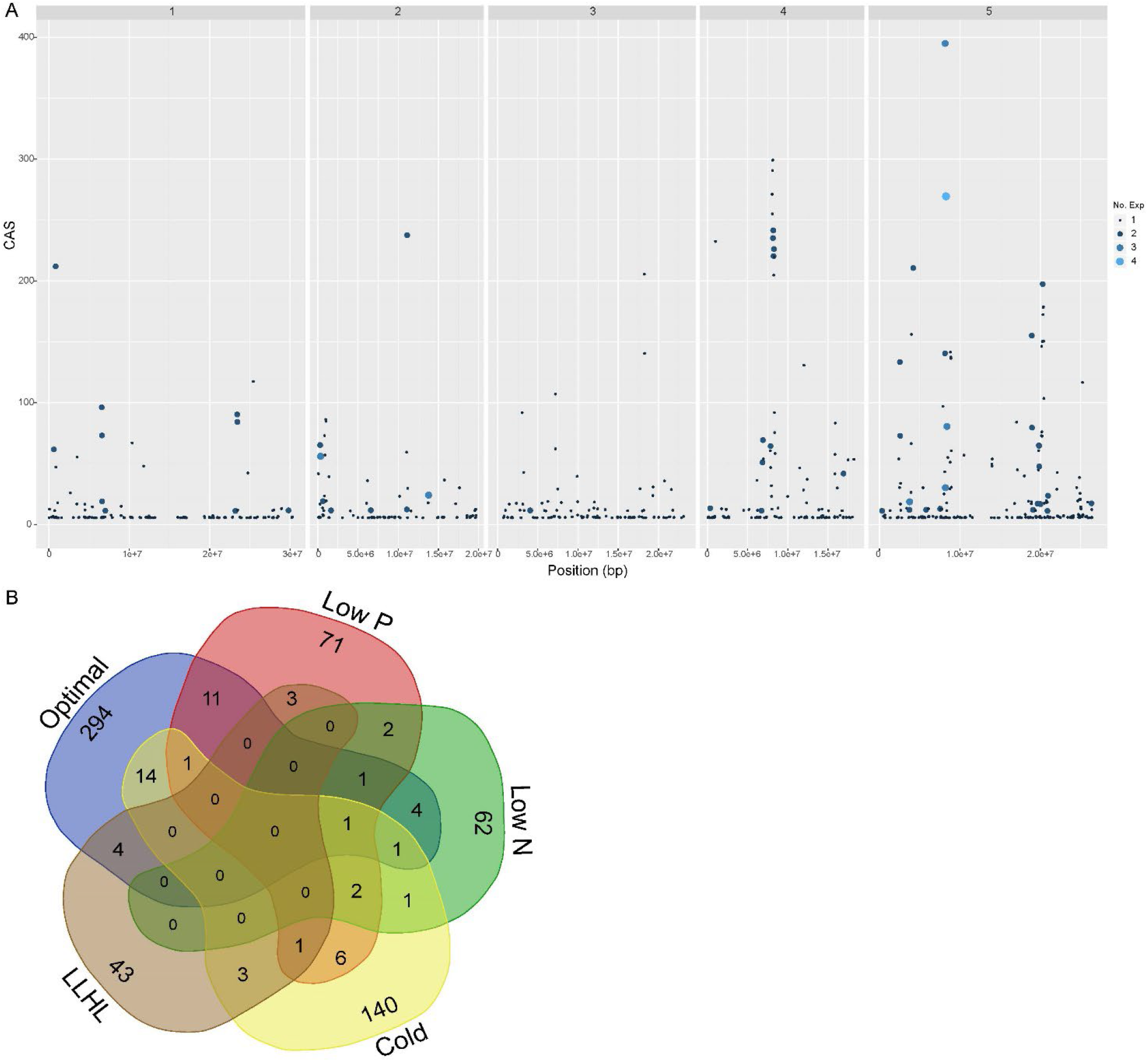
Comparison of genetic loci for photosynthetic light-use efficiency in multiple experiments. (A) Significant QTLs with individual −log_10_(p) ≥ 5.5 in every single GWAS analysis throughout the five datasets generated from five independent experiments were integrated and visualized into a plot. Each significant QTL with a genomic window of 25 kb is presented as a dot at its physical position along the five chromosomes of *A. thaliana* (x-axis) and the its cumulative association score (CAS, y-axis). The CAS is the sum of significant QTL −log_10_(p) values obtained in individual GWAS, over the total of 332 analyses. The size and color of the dot indicate in how many independent experiments (No. Exp) the QTL was identified, ranging from 1 to 4. (B) The composition of the QTL inventory is shown in a Venn diagram, which is attributed to Optimal, Low phosphorus supply (Low P), Low nitrogen supply (Low N), changing temperature and darkness treatment (in short Cold) and changing irradiance (low light to high light, LLHL) experiment.

**Table 1.**
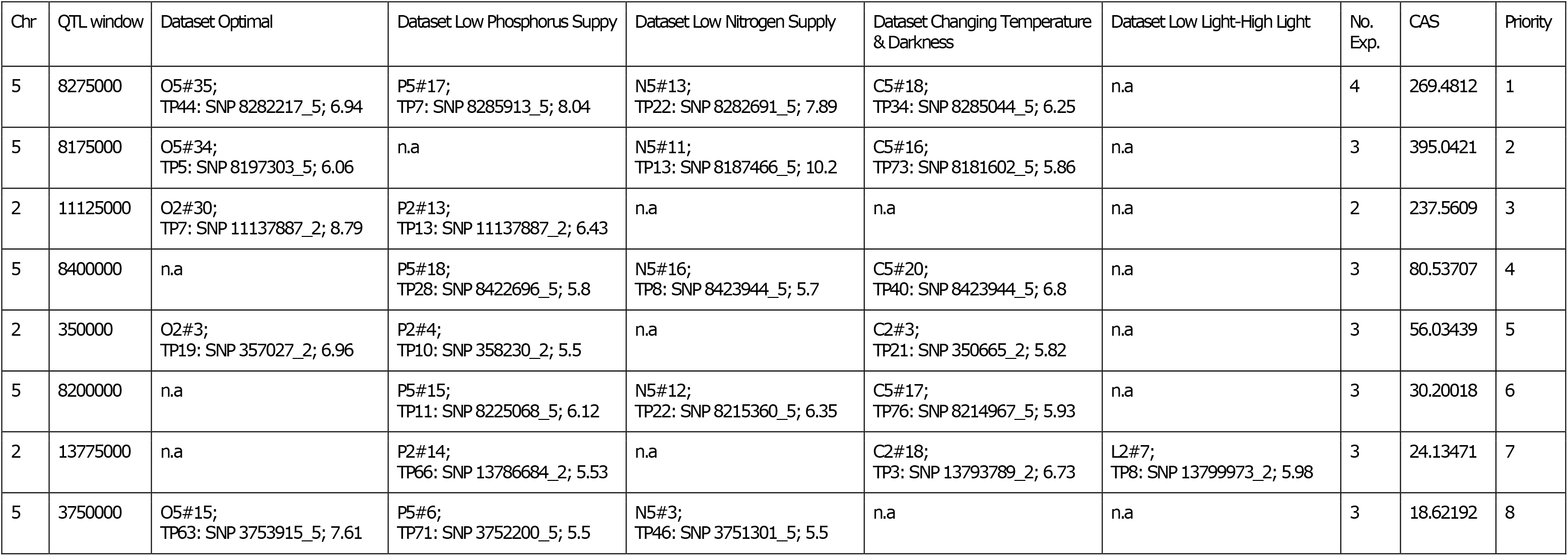
Details of top eight QTLs identified in this study. The top QTLs are reported for their location on chromosome (Chr.), QTL window, dataset in which the QTL is found, the total number of experiments in which the QTL is detected, the cumulative association score (CAS) and the priority rank. The CAS is the sum of significant QTL −log_10_(p) values obtained in individual GWAS, over the total of 332 analyses. A representative timepoint (TP) of each dataset is selected where the QTL is most significant, which contains the most significant SNP associated with phenotypic variation and its respective −log10(p) score. The total number of experiments in which the QTL is detected is shown (No. Exp.). Not detected QTL is indicated by ND

| Chr | QTL window | Dataset Optimal | Dataset Low Phosphorus Supply | Dataset Low Nitrogen Supply | Dataset Changing Temperature & Darkness | Dataset Low Light-High Light | No. Exp. | CAS | Priority |
| --- | --- | --- | --- | --- | --- | --- | --- | --- | --- |
| 5 | 8275000 | O5#35;<br>TP44: SNP 8282217_5; 6.94 | P5#17;<br>TP7: SNP 8285913_5; 8.04 | N5#13;<br>TP22: SNP 8282691_5; 7.89 | C5#18;<br>TP34: SNP 8285044_5; 6.25 | n.a | 4 | 269.4812 | 1 |
| 5 | 8175000 | O5#34;<br>TP5: SNP 8197303_5; 6.06 | n.a | N5#11;<br>TP13: SNP 8187466_5; 10.2 | C5#16;<br>TP73: SNP 8181602_5; 5.86 | n.a | 3 | 395.0421 | 2 |
| 2 | 11125000 | O2#30;<br>TP7: SNP 11137887_2; 8.79 | P2#13;<br>TP13: SNP 11137887_2; 6.43 | n.a | n.a | n.a | 2 | 237.5609 | 3 |
| 5 | 8400000 | n.a | P5#18;<br>TP28: SNP 8422696_5; 5.8 | N5#16;<br>TP8: SNP 8423944_5; 5.7 | C5#20;<br>TP40: SNP 8423944_5; 6.8 | n.a | 3 | 80.53707 | 4 |
| 2 | 350000 | O2#3;<br>TP19: SNP 357027_2; 6.96 | P2#4;<br>TP10: SNP 358230_2; 5.5 | n.a | C2#3;<br>TP21: SNP 350665_2; 5.82 | n.a | 3 | 56.03439 | 5 |
| 5 | 8200000 | n.a | P5#15;<br>TP11: SNP 8225068_5; 6.12 | N5#12;<br>TP22: SNP 8215360_5; 6.35 | C5#17;<br>TP76: SNP 8214967_5; 5.93 | n.a | 3 | 30.20018 | 6 |
| 2 | 13775000 | n.a | P2#14;<br>TP66: SNP 13786684_2; 5.53 | n.a | C2#18;<br>TP3: SNP 13793789_2; 6.73 | L2#7;<br>TP8: SNP 13799973_2; 5.98 | 3 | 24.13471 | 7 |
| 5 | 3750000 | O5#15;<br>TP63: SNP 3753915_5; 7.61 | P5#6;<br>TP71: SNP 3752200_5; 5.5 | N5#3;<br>TP46: SNP 3751301_5; 5.5 | n.a | n.a | 3 | 18.62192 | 8 |

From this QTL inventory, we prioritized the top eight QTLs according to their reproducibility in multiple environments and their CAS (Table 1) for further LD analysis to identify candidate genes (Supplemental table 7). The LD regions around these QTLs comprised between two to 12 candidate genes. None of the candidate genes has been previously reported to be associated with variation in photosynthesis efficiency, except for the QTL on chromosome 5 at position 8175000 bp. This particular QTL, which is detected in four conditions, has previously been identified to be associated with natural genetic variation at a cluster of *SQUALENE MONOOXYGENASE* (*SQE*)-like genes (*SQE5-SQE7*) in recombinant inbred line population between Ler-0 and Col-0 in three conditions (optimal, low light and low nitrogen supply) (van Bezouw et al., 2023). Altogether, it suggests that SQE QTL is a very robust to affect photosynthesis in diverse environments.

## Discussion

We assessed photosynthetic light-use efficiency, determined as ΦPSII, at the growth or treatment irradiance in the HapMap diversity set of *A. thaliana* accessions. These were phenotyped in five independent experiments in the high-throughput phenotyping platform in Wageningen University (Flood et al., 2016). Next to the ‘optimal’, (i.e. non-stressful) conditions, plants were exposed to several stress-inducing conditions, such as low N or P supply, low growth temperatures or a single-step increase in irradiance. The large datasets that were obtained allowed the examination of the ΦPSII response to adverse conditions and the identification of QTLs corresponding to natural genetic allelic variation associated with the phenotypic response to these environments.

Our study revealed that there is a large genetic variation for ΦPSII (>600 loci with very small effect, figure 6) despite the relatively low broad sense heritability observed for this trait (Supplemental Figure 1). This fact is underlined by the complex polygenic nature of photosynthesis and high phenotypic dynamic because of tightly regulating processes at biochemical level and strongly influence of micro-environment.

The dynamic nature of photosynthesis is also reflected by the detection of many QTLs unique for either a specific growth phase or time of the day in this study. In steady conditions (optimal, low N and low P), changes of associated QTL profiles are observed that coincided with two changes in growth, around 16 DAS and 20 DAS (if the experiments lasted longer than 24 DAS, in control and low P experiment, Fig 1, 4 and 7). These changes probably mark the divergence in growth among genotypes, which has been estimated to be occurring at the end of the lag phase (16 DAS) and the beginning of the exponential growth phase (20 DAS), as determined by the rosette diameter or the effective rosette leaf surface area (Wieters et al., 2021, Gonzalez et al., 2020).

Several loci are found to be rhythmic, and therefore suggested to be affected by the circadian clock, or the diurnal rhythm. Diurnal fluctuations in Chl a/b ratio, displaying a contrasting phase when compared to the rate of biosynthesis and degradation of the PSII light-harvesting complex, have been interpreted as drivers of the circadian rhythm in photosynthesis in bean and cotton (García-Plazaola et al., 2017). The dip in Chl a/b ratio occurs at noon, which coincides with the maximal expression of PSII light harvesting complex protein levels. This supports the observation that ΦPSII in constant condition is higher at noon than at the afternoon timepoint.

The identified QTLs for photosynthesis in our study disperses across the genome and with a very small effect each, which is consistent with an omnigenic architecture, as originally proposed for human disease response (Boyle et al. 2017) and later adopted for crop yield (Hu et al., 2024). Omnigenic model describes a limited number of core genes that are embedded within extensive regulatory networks of peripheral (pleiotropic) genes. This network-level control may be even more pronounced due to tight metabolic regulation. In addition to that photosynthesis is likely optimized by natural selection (Denison, 2012), for example Rubisco was described as almost perfectly optimal (Tcherkez et al., 2006). Therefore in our QTL inventory, the chance to identify core genes with large effect is unlikely but rather predominant peripheral genes with small effects.

Out of 655 QTLs, only for a few QTLs the causal gene(s) was verified. A shared QTLs for PSII in response to cold and an increase in irradiance on chromosome 5 window 4200000 appears to cause by a allelic variation of the *DGS1* (*DIGALACTOLIPID-DEFICIENT MUTANT 1 SUPPRESSOR 1*) gene, as previously confirmed by phenotypic analysis of a *dgs1* T-DNA knock-out mutant in response to cold and an increase in irradiance (van Rooijen et al., 2017, Prinzenberg et al., 2020). *DGS1* is important for the conversion of phospholipids to galactolipids and induced in response to high-light stress (van Rooijen et al., 2018). Another common QTL, located on chromosome 1 window 850000, and identified under optimal conditions and upon cold stress is caused by the variation at the *PSB27* (Prinzenberg et al., 2020) gene encoding a thylakoid lumen protein associated with the PSII complex and involved in the repair and maintenance of PSII complex proteins. Copy number variation of *SQE* genes (*SQE5-SQE7*), of which the molecular function is still unknown, are responsible for photosynthetic variation at QTL on chromosome 5 window 817500, that was identified in the optimal, low N and changing temperature in our study and also in low light condition in the study of van Bezouw et al. (2023).It is notable that these verified QTLs leading to causal gene discovery are robust QTLs having effect in multiple environments. It is still very challenging to discover gene underlying QTLs with specificity and very small effect. In these three successful cases, causal genes do not directly control the regulation of photosynthesis processes. This exuberates the observation that natural genetic variation for photosynthesis predominantly is expressed via peripheral genes.

In breeding for a complex trait like crop yield, the factors affecting the trait can be categorised into: (1) physiological adaptive traits (e.g. stomatal control, leaf growth, sensitivity of leaf expansion, stress responses) that vary several-fold over minutes to hours as response to environmental condition; (2) constitutive traits (e.g. plant architecture traits, flowering time, reproductive development) that show more long-term variation (Welcker et al. 2022). Welcker et al. (2022) showed that stable maize yield in different environmental conditions were selected based on the constitutive traits, and physiological adaptive traits only give advantage in certain circumstances and therefore not selected by breeder. Similar to crop yield, photosynthesis are affected by many factors including both physiological adaptive traits and constitutive traits, which implicates that robust QTLs in multiple conditions for photosynthesis in our study are likely underlined by constitutive-trait driving genes than those drive physiological adaptive traits. These specific QTLs regulating physiological adaptive traits are an allele reservoir potentially under challenging conditions.

Numerous studies have demonstrated substantial natural variation in photosynthetic traits among genotypes in major crop species, including rice (Qu et al., 2017), wheat (Driver et al., 2014; Carmo-Silva et al., 2017), soybean (Wang et al., 2020) indicating that photosynthetic performance has not been fully optimized through breeding. Although these studies largely describe phenotypic variation without resolving its underlying genetic basis, the extent of variation is remarkably similar to that observed in *A. thaliana*. Our results show that this phenotypic variation in *Arabidopsis* is not explained by a few major loci, but instead by a large number of QTLs distributed throughout the genome. This suggests that the extensive phenotypic variation observed in crop species is also likely to arise from a similarly polygenic genetic architecture. Consequently, many of the *A. thaliana* genes underlying ΦPSII QTLs could potentially be investigated as a target for marker assisted breeding for increased photosynthetic efficiency in crops. However, given the large number of loci involved with small effect size it seems unlikely that a marker assisted breeding approach will contribute to a fast increase of photosynthesis efficiency in crop and potentially a higher crop biomass or yield. Instead, breeding approaches that capture the cumulative effects of many small-effect loci, such as genomic prediction, are likely to be more effective (Sharma et al., 2021; Zaim et al., 2020). In addition, the omnigenic model has become a new quantitative genomics framework in plant system, where the integration of classical quantitative genetics with molecular and developmental genetics may be unified to understand better the genetic of complex traits such as photosynthesis.

## Methods

### Plant material

In this work, we have used 633 *A. thaliana* accessions belonging to two diversity panels of natural accessions, the Hapmap panel (350 genotypes) of global accessions (Atwell et al., 2010; Baxter et al., 2010; Li et al., 2010) and the Swedish RegMap panel (283 genotypes) (Horton et al., 2012). The Hapmap panel was screened for photosystem II efficiency (ΦPSII) in all five experiments including (1) optimal nutrient supply (Optimal), (2) low phosphorus supply (Low P), (3) low nitrogen nutrient supply (Low N), (4) fluctuating temperature (from 21°C to 5°C) (Prinzenberg et al., 2020) followed by acclimation back to the initial condition (20°C) after a period of darkness (in short, called ‘Cold’ treatment) and (5) a one-step increase in irradiance (from low light to high light, or ‘LLHL’ treatment) (van Rooijen et al., 2017) (Supplemental Table 8). The Swedish RegMap panel was only tested in optimal conditions.

### Experimental design and photosynthesis measurements

In general, plants were grown in a climate-controlled growth chamber on rockwool blocks (Grodan, Rockwool Group, The Netherlands, 40 × 40 × 40 mm in size) supplied hydroponically with a nutrition solution and at controlled settings for photoperiod, light intensity during the day and day/night temperature (Supplemental Table 7). Relative humidity was set at 70%. Unless specified, standard environmental parameters were applied. Standard nutrition solution was composed of macronutrients at the following final elemental composition of dissolved ions: NH_4_^+^ 1.7 mM, NO_3_^-^ 4.14 mM, PO ^-/2-^ 1.29 mM, K^+^ 4.13 mM, Ca^2+^ 1.97 mM, Mg^2+^ 1.24 mM, SO ^2-^ 3.14 mM, Fe^2+^ 21 μM (composed half-half of Fe-DTPA and Fe-EDDHA), Mn^2+^ 3.4 μM, Zn^2+^ 4.7 μM, BO_3_^3-^ 14 μM, Cu^2+^ 6.9 μM, MoO ^2-^ 0.5 μM. The standard solution was adjusted with KOH or H_2_SO_4_ to a pH of 6.2, the final EC was 1.4. Standard light irradiance (PAR) was set at 200 μmol m^-2^ s^1^ in combination with a standard photoperiod of 10/14 h day/night. The day/night temperature was set at 20/18°C. Every accession was grown in four replicates in all experiments, except for the control experiment.

Photosynthesis was measured as ΦPSII, the light-use efficiency of photosystem II electron transport, based on chlorophyll fluorescence images of plants using a high-throughput phenotyping system as described by Flood et al. (2016). Images were taken at multiple times per day during the course of the experiment, which resulted in a total of 218 datapoints from the five above mentioned experiments.

The specific environmental settings and ΦPSII data collection for each experiment (Table 1) are described as follows:

1. The ‘Optimal’ experiment was conducted over six sub-experiments that assured a minimum of four replicates and a maximum of eight replicates for each accession. ΦPSII was measured three times a day from 11 days after sowing (DAS) to 24 DAS, which gives a total of 41 data- timepoints.
2. In the ‘Low P’ experiment, plants were grown under low phosphorus supply of 0.1 mM, as compared to 1.29 mM in the standard nutrition solution. Photosynthesis was measured three times a day from 10 to 26 DAS leading to a total of 51 data-timepoints.
3. The ‘Low N’ experiment was conducted using a modified standard nutrient solution, containing 0.1 mM NH_4_^+^ and 0.904 mM NO_3_^-^, which equals 17% of total N available in the standard solution (1.7 mM NH_4_^+^ and 4.14 mM NO_3_^-^). Chlorophyll fluorescence images were taken from 12 to 32 DAS for three times a day, which makes 32 ΦPSII data-timepoints.
4. In the ‘Cold’ experiment, both the effect of low temperature (Part 1) and of a period of darkness (Part 2) on ΦPSII response were examined. The first part of this experiment, that was published by Prinzenberg et al. (2020), covered the first 22 DAS. In brief, plants were grown at regular temperature (21°C) until the end of 13 DAS, then exposed to a period of seven days at low temperature (5°C) (from 14 DAS to the end of 20 DAS), and allowed to recover for two days at regular temperature (21°C). The second part started from 23 DAS until the end of the experiment, 33 DAS, in which a period of complete darkness was given to plants for four days (from 23 DAS to the end of 26 DAS) and standard light conditions were applied thereafter (from 27 DAS to 33 DAS). Different from the four other experiments, the photoperiod was set at 12/12 h day/night. The ΦPSII was measured four times a day from 11 DAS in the first part and from 27 DAS in the second part, resulting in 76 data-timepoints.
5. Van Rooijen et al. (2017) previously described the ‘LLHL’ experiment in detail. Briefly, plants were grown under low light conditions of 100 μmol m^-2^ s^-1^ for the first 24 days, thereafter the irradiance was increased to 550 μmol m^-2^ s^-1^. ΦPSII was measured three times a day from 23 till 28 DAS, resulting in 18 data-timepoints.

### Statistical analyses

#### Phenotypic analysis

The raw chlorophyll fluorescence images were converted into average ΦPSII values per plant, as a phenotypic parameter of photosynthesis (Flood et al., 2016). The phenotypic mean of the replicates was calculated as the best linear unbiased estimator (BLUE) using a linear mixed model adjusted for experimental design factors (Flood et al., 2016). In most of the experiments, important factors are spatial row (x) and column (y) coordinates, image position and sowing block. Only for the control experiment of the HapMap and the RegMap, the mean was adjusted for sub-experiment factor instead of sowing block factor. The equation below was implemented in R with the lmer function and lme4 package.

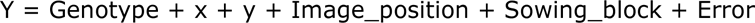

where Genotype is used as fixed effect and the others factors are defined as random effects.

#### Heritability estimates

The broad sense heritability was estimated as the ratio of estimated genetic variance over the sum of estimated genetic and environmental variance described in the phenotypic analysis. These variances were calculated using the equation above with the REML function and implemented in R.

### Genome-wide association analysis and data visualization

#### Genome-wide association analysis

GWAS were performed using univariate and multivariate linear mixed models implemented in the GEMMA software (Zhou and Stephens, 2012). The Univariate linear mixed model (uLMM) is fitted for marker association tests with a single phenotype to account for population structure (Zhou and Stephens, 2012). In the multivariate linear mixed model (mLMM), markers are tested for association with multiple phenotypes simultaneously, while controlling for population structure (Zhou and Stephens, 2017). We have used the BLUEs of ΦPSII at single datapoint as input phenotypes for GWAS. This analysis was performed for the HapMap population using the 1 M imputed SNPs genotype dataset (Arouisse et al., 2020). Rare SNPs with a minor allele frequency below 0.05 were removed and population structure was accounted for in each analysis.

The statistical significance threshold of the association test was determined at the adjusted Bonferroni levels. The adjusted Bonferroni threshold was based on the assumption that SNPs are not segregating independently, but in association with other SNPs in linkage disequilibrium (LD). Kim et al. (2007) reported that LD decays rapidly on average within 10 kb in natural population, meaning roughly 15000 LD blocks in the *A. thaliana* genome. Therefore we estimated the adjusted Bonferroni threshold of −log_10_(0.05/150000), equivalent to 5.5 and used this as significance threshold in our analyses to report marker-trait associations at QTLs.

#### The synthesized QTL inventory

To integrate and visualize results of all GWAS into one QTL inventory, we developed a ‘Turbo mapper’ in R, that divides the genome into 25-kb window and reports the −log_10_(p) score of the window (if ≥ 5.5 threshold) based on the most significant SNP in the window. In the next step, all output results are scanned to sum up the window −log_10_(p) value into a cumulative association score (CAS) and report the count of how many times the QTL windows were significant throughout the total number of individual analyses. The CAS, that indicates how significant and how often the window is detected as a QTL, is visualized in the Manhattan plot for every window throughout the five chromosomes. This mapping tool also provides a Signal file that contains further details (window position, individual −log_10_(p) value in individual analysis, count and CAS of the reported QTL window.

#### Linkage disequilibrium analysis (LD) and candidate gene identification

The top eight prioritized QTLs (Table 1) were further investigated with LD analysis to determine QTL intervals and as a result the identification of candidate genes. QTLs were searched separately in each dataset in which they were found to identify the most significant SNP as a target SNP for LD analysis (Supplemental Table 7). For QTLs found in multiple datasets, the most significant SNP can be different, and all of them were considered target SNP. The correlation between other SNPs in the surrounding genomic region of 50 kb and the target SNP was calculated using the gastron package in R. A cutoff threshold of 0.4 was used to determine the correlation and thus LD region. All genes which reside within the LD region were listed as candidate genes (Supplemental Table 7).

## Author contributions

JH and MGMA conceived the project. PJF, CNM and NOE designed and performed the plant screening for photosynthesis experiments. T-PN analysed the data. TPJMT contributed to data analyses and visualization. T-PN wrote the manuscript with contributions from MGMA and JH. All authors read and commented on the manuscript.

## Acknowledgements

Plants of the experiments were taken care by Gerrit Stunnenberg and Taede Stoker of Unifarm, to whom we are grateful. During this project, there were a number of students whose support is acknowledged: Ioanis Thannos, Siyuan Wei, Sumit Singh and Jacky To. We like to thank Dr. Roel van Bezouw for his practical help, discussions and suggestions about data analysis and interpretation. The data of the second part in the cold experiment (after darkness treatment) was kindly provided by Dr. Aina E. Prinzenberg, for which we are grateful.

## Supplemental information

**Supplemental Table 1**. Excel file detailed QTL inventory for Optimal condition

**Supplemental Table 2**. Excel file detailed QTL inventory for low phosphorus supply condition

**Supplemental Table 3**. Excel file detailed QTL inventory for low nitrogen supply condition

**Supplemental Table 4**. Excel file detailed QTL inventory for changing temperature and darkness condition

**Supplemental Table 5**. Excel file detailed QTL inventory for step change in irradiance low light to high light condition

**Supplemental Table 6**. Excel file synthesized QTL inventory for all five datasets

**Supplemental Table 7.** Excel file analysis of LD and candidate genes for top eight QTLs

**Supplemental Table 8.** Overview of environmental conditions for the five experiments used in this work. The five experiments listed are optimal nutrient supply (Optimal), low phosphorus supply (Low P), low nitrogen supply (Low N), changing temperature followed by acclimation after a period of darkness (Cold) and a one-step increase in irradiance (LL-HL). The types of population (Population) and number of individuals (No. Individuals) evaluated in the experiment are specified. Environmental variables for each experiment are given as Photo period (hours of day/night), light intensity (Irradiance, μmol.m^-2^.s^-1^),and Temperature (°C) during day and night. Experimental setup includes the number of replicates (No. Rep), the period (Start-End) of days after sowing (DAS) during which photosystem II efficiency (ΦPSII) was measured, the number of ΦPSII datapoints per day and the resulting total number of data timepoints (Total no. timepoints) in the experiment.

| Experiment | Population | No. Individuals | Photo period (day/night) hr | Irradiance $\mu\text{mol.m}^{-2}.\text{s}^{-1}$ | Temperature (day/night) $^{\circ}\text{C}$ | No. Rep | $\Phi\text{PSII}$ measurement (Start - End) DAS | $\Phi\text{PSII}$ datapoints/day | Total no. timepoints | Publication |
| --- | --- | --- | --- | --- | --- | --- | --- | --- | --- | --- |
| (1) Optimal | HapMap & RegMap | 633 | 10/14 | 200 | 20/18 | 8 | 11 – 24 | 3 | 41 | NA |
| (2) Low P | HapMap | 350 | 12/12 | 200 | 20/18 | 4 | 10 - 26 | 3 | 51 | NA |
| (3) Low N | HapMap | 350 | 10/14 | 200 | 20/18 | 4 | 12 – 21 | 3 | 32 | NA |
| (4) Cold | HapMap | 350 | 12/12 | 200 | 21/18 - 5/5 - 21/18 | 4 | 11 – 22 & 27 – 33 | 4 | 76 | Prinzenberg et al., 2020, and NA |
| (5) LLHL | HapMap | 350 | 10/14 | 100 - 500 | 20/18 | 4 | 23 – 28 | 3 | 18 | Van Rooijen et al., 2017 |

**Supplemental Figure 1.**
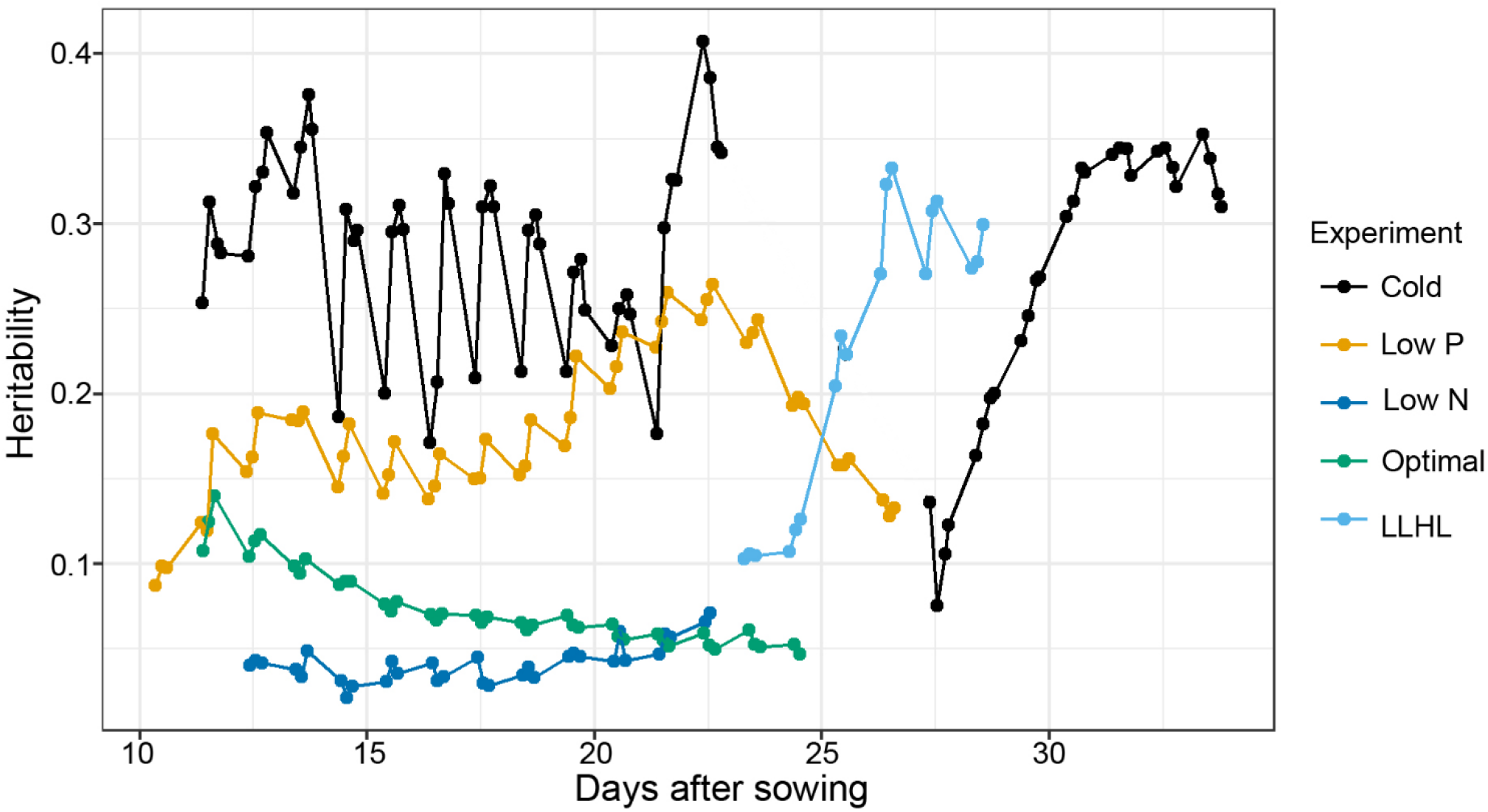
Broad-sense heritability for ΦPSII in the five experiments. Broad sense heritability (H^2^; y-axis) was calculated for ΦPSII at each timepoint (indicated as filled circle) over the course of five experiments, as indicated in ‘days after sowing’ (x-axis). The colours represent independent experiments: green for optimal, yellow for low phosphorus supply (Low P), dark blue for low nitrogen supply (Low N), black for fluctuating temperature (from normal to cold temperature) followed by acclimation to normal condition after a period of darkness (in short, Cold), and light blue for fluctuating irradiance (low light to high light, LLHL).

**Supplemental Figure 2.**
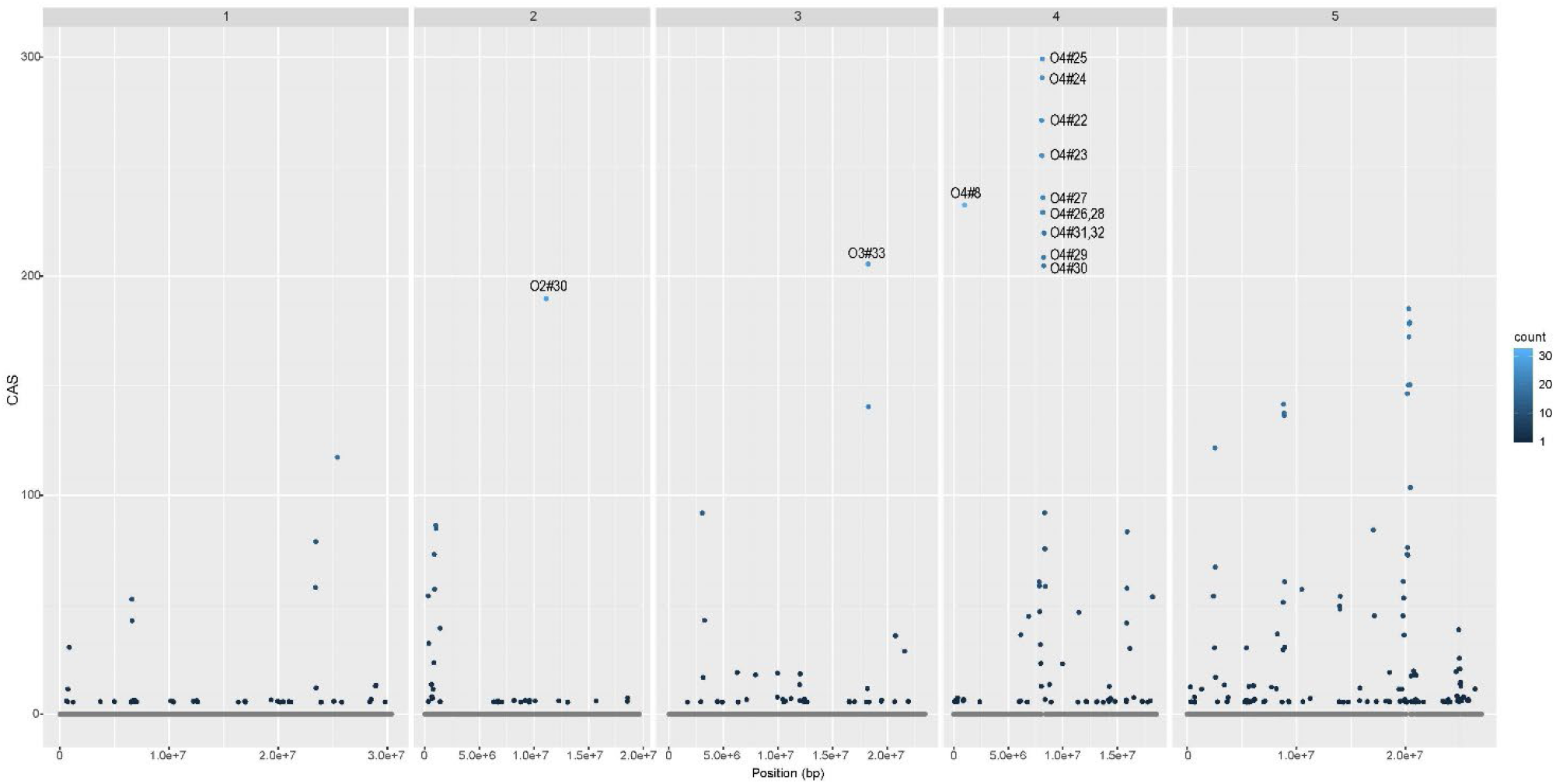
QTL inventory for photosynthesis under optimal conditions. Significant QTLs with individual −log_10_(p) ≥ 5.5 in every single GWAS analysis throughout the dataset generated in optimal experiment were combined and visualized into a plot. Each significant QTL, corresponding to a genome window of 25 kb, is presented as a dot based on its physical position along the five chromosomes of Arabidopsis (x-axis) and the its cumulative association score (CAS, y-axis). The CAS is the sum of significant QTL −log_10_(p) values in individual GWAS over the total of 64 analyses. The color of each dot indicates the number of times (count) it has been identified as significant in an individual analysis, ranging from 1 to 32. QTLs with the highest CAS are labelled.

**Supplemental Figure 3.**
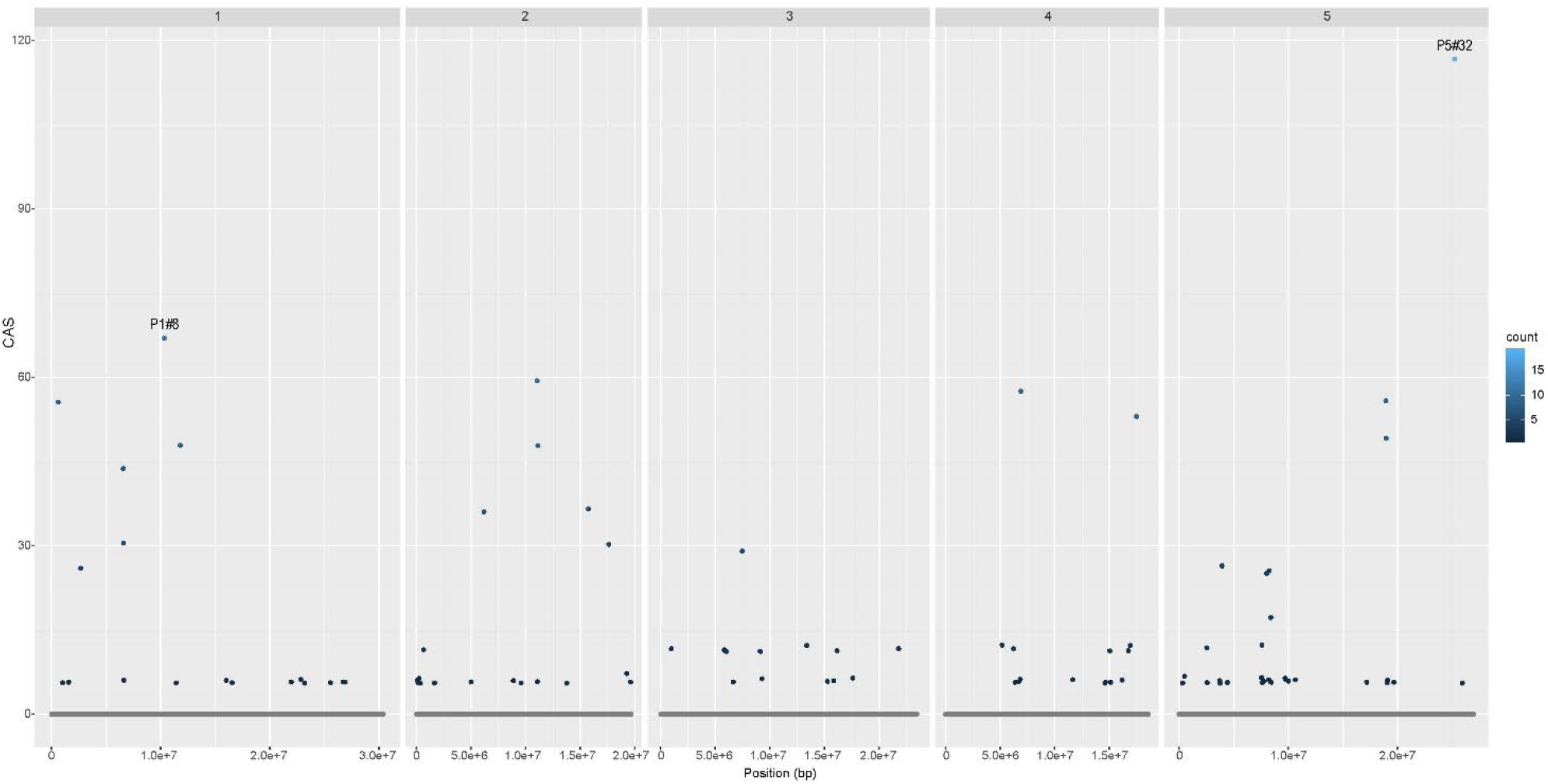
QTL inventory for photosynthesis under low phosphorus supply condition. Significant QTLs with individual −log_10_(p) ≥ 5.5 in every single GWAS analysis throughout the dataset generated in low phosphorus supply experiment were combined and visualized into a plot. Each significant QTL, corresponding to a genome window of 25 kb, is presented as a dot based on its physical position along the five chromosomes of Arabidopsis (x-axis) and the its cumulative association score (CAS, y-axis). The CAS is the sum of significant QTL −log_10_(p) values in individual GWAS over the total of 77 analyses. The color of each dot indicates the number of times (count) it has been identified as significant in an individual analysis, ranging from 1 to 19. QTLs with the highest CAS are labelled.

**Supplemental Figure 4.**
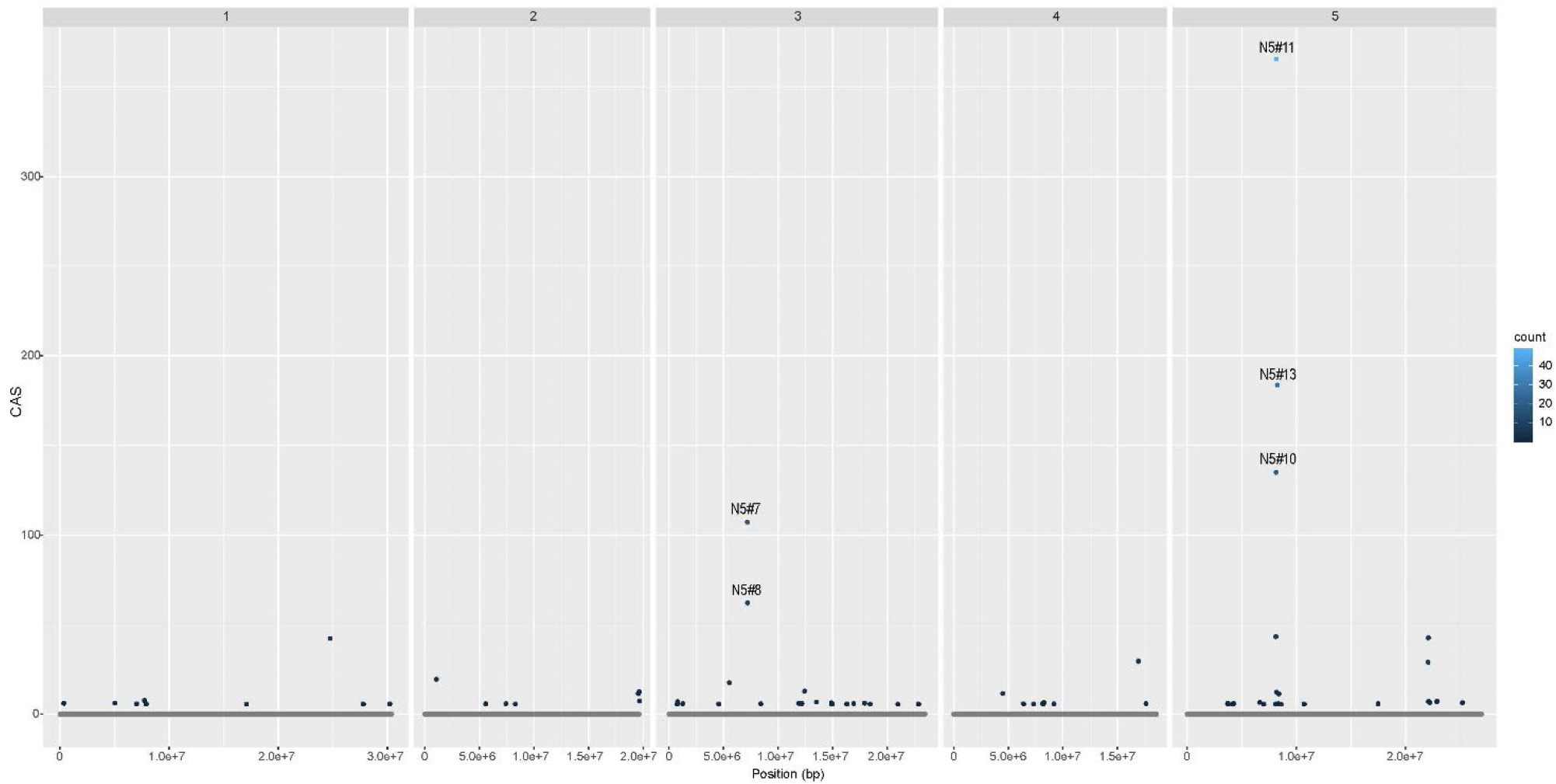
QTL inventory for photosynthesis under low nitrogen supply condition. Significant QTLs with individual −log_10_(p) ≥ 5.5 in every single GWAS analysis throughout the dataset generated in low nitrogen supply experiment were combined and visualized into a plot. Each significant QTL, corresponding to a genome window of 25 kb, is presented as a dot based on its physical position along the five chromosomes of Arabidopsis (x-axis) and the its cumulative association score (CAS, y-axis). The CAS is the sum of significant QTL −log_10_(p) values in individual GWAS over the total of 49 analyses. The color of each dot indicates the number of times (count) it has been identified as significant in an individual analysis, ranging from 1 to 49. QTLs with the highest CAS are labelled.

**Supplemental Figure 5.**
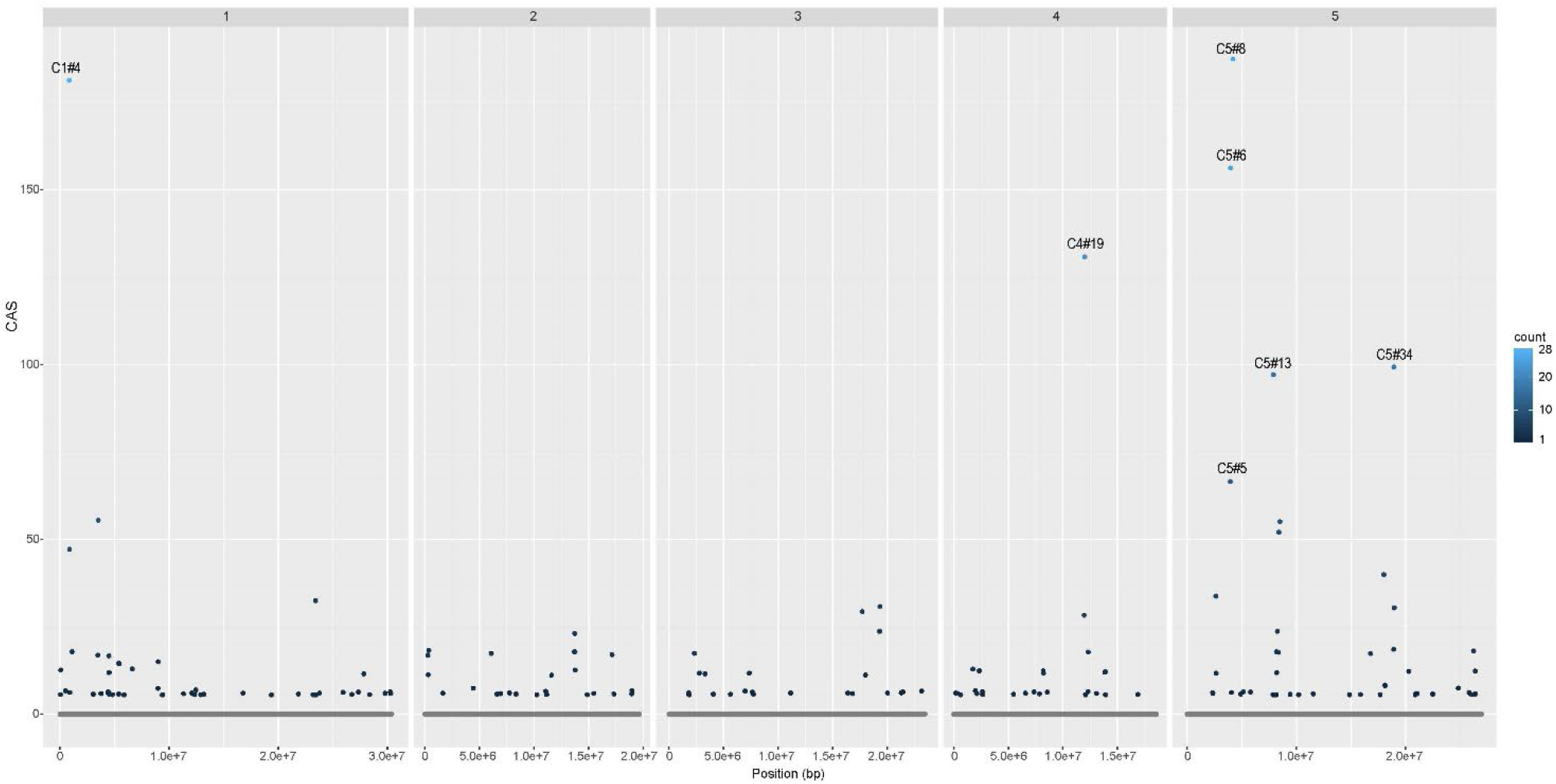
QTL inventory for photosynthesis under fluctuating temperature condition and after darkness treatment. Significant QTLs with individual −log_10_(p) ≥ 5.5 in every single GWAS analysis throughout the dataset generated in fluctuating temperature and after darkness treatment experiment were combined and visualized into a plot. Each significant QTL, corresponding to a genome window of 25 kb, is presented as a dot based on its physical position along the five chromosomes of Arabidopsis (x-axis) and the its cumulative association score (CAS, y-axis). The CAS is the sum of significant QTL −log_10_(p) values in individual GWAS over the total of 111 analyses. The color of each dot indicates the number of times (count) it has been identified as significant in an individual analysis, ranging from 1 to 28. QTLs with the highest CAS are labelled.

**Supplemental Figure 6.**
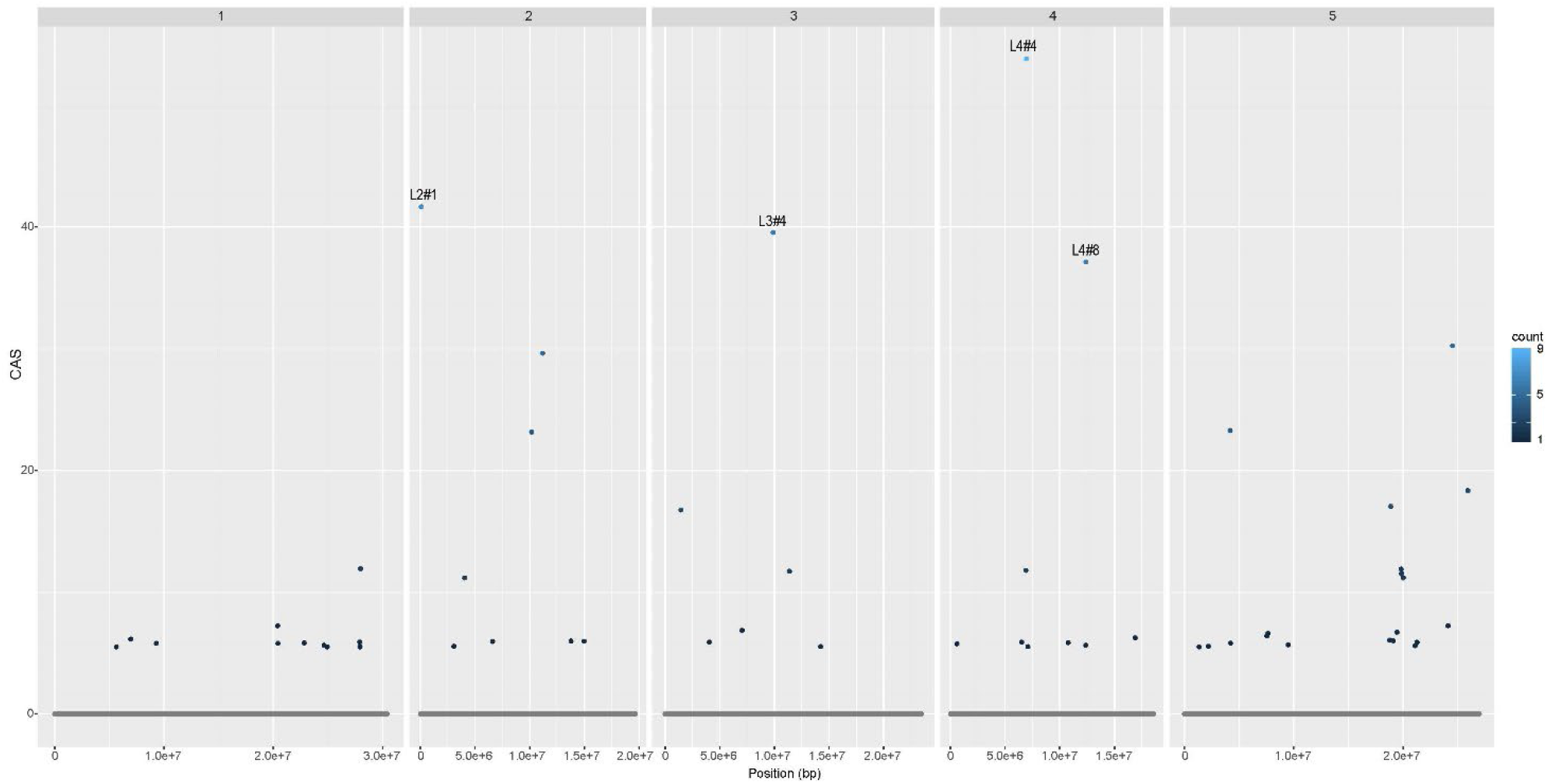
QTL inventory for photosynthesis under changing irradiance (low light to high light). Significant QTLs with individual −log_10_(p) ≥ 5.5 in every single GWAS analysis throughout the dataset generated in changing irradiance (low light to high light) experiment were combined and visualized into a plot. Each significant QTL, corresponding to a genome window of 25 kb, is presented as a dot based on its physical position along the five chromosomes of Arabidopsis (x-axis) and the its cumulative association score (CAS, y-axis). The CAS is the sum of significant QTL −log_10_(p) values in individual GWAS over the total of 30 analyses. The color of each dot indicates the number of times (count) it has been identified as significant in an individual analysis, ranging from 1 to 9. QTLs with the highest CAS are labelled.

